# Convergent routes to haloterpenoid biosynthesis in red algae: identification of non-canonical UbiA-type terpene synthase gene clusters in the genus *Laurencia*

**DOI:** 10.64898/2026.10.28.762581

**Authors:** Hannah K. Bone, Aodhan W. Beattie, Sheila Podell, Ashleigh J. Burke, Taylor S. Steele, Immo Burkhardt, Bradley S. Moore

**Author notes:** Bradley S. Moore, **Email:**.

## Abstract

Red algae produce structurally diverse haloterpenoids unique to seaweeds, yet the enzymes responsible for generating many of their core terpene scaffolds remain unknown. Here, we report the identification of red algal UbiA-type terpene synthases (UTSs), a noncanonical enzyme family recently associated with terpene biosynthesis in bacteria and fungi. By prioritizing UbiA candidates located near putative biosynthetic genes, we identified eight candidate gene clusters across two red algal species, *Laurencia pacifica* and *Laurencia subopposita*. Using the heterologous host plant *Nicotiana benthamiana*, we show that the UbiA genes act as non-canonical TSs and that two UTSs produce the sesquiterpene bisabolene. The discovery of the bisabolene UTSs, LsUTS-2 and LpUTS-2, lays the foundation for laurane and chamigrane-type sesquiterpenoid production, addressing a prior biosynthetic quandary that bisabolene is not produced by established red algal class I TSs. Phylogenetic analyses distinguish the red algal UTSs from canonical UbiA prenyltransferases involved in primary metabolism and suggest they were acquired outside the red algal lineage. These findings expand the known distribution of UTSs and establish a previously unrecognized route to red algal terpene scaffolds.

## Introduction

Terpenoids represent the largest and most structurally diverse class of natural products. This diversity arises from the cyclization and rearrangement of linear isoprenoid precursors into a wide range of core scaffolds, which can then be further modified by tailoring enzymes. Marine organisms are further able to expand this diversity through the enzymatic incorporation of halide ions, which are present in the ocean environment, to generate unique haloterpenoids. Red algae (Rhodophyta), particularly members of the genus *Laurencia*, are one of the most prolific producers of haloterpenoids.^1^ Often, these organohalogens feature patterns of mixed halogenation in which both chlorine and bromine atoms are incorporated into a single product, thereby representing a distinct hallmark of red algal metabolites.^2^ Although numerous terpenoids with distinct scaffolds have been isolated from *Laurencia*, chamigrane- and laurane-type sesquiterpenoids are among the most prevalent.^3^ These molecules often play important ecological roles, as exemplified by the chamigrane terpenoid, elatol, which has been demonstrated to deter predation and act as a chemical barrier to protect the alga from biofouling.^4,5^ Their biological relevance, alongside growing interest in halogenated metabolites for therapeutic development, has long motivated researchers to identify the biosynthetic pathways that red algae use to produce these structurally complex molecules.^6,7^

Terpene scaffold formation is commonly initiated by class I terpene synthases (TSs), which ionize prenyl diphosphate substrates to generate reactive carbocation intermediates. These intermediates undergo tightly controlled cyclization and rearrangement reactions within the enzyme active site, producing either a single scaffold or a mixture of products.^8^ Enzymes of this type could be identified in *Plocamium pacificum* and *Portieria hornemannii*, where they were responsible for producing the monoterpenes ocimene and myrcene, the precursors for hallmark halogenated monoterpenoids like halomon found in these red algal species. Previous work in two *Laurencia* species*, L. pacifica* and *L. subopposita*, also characterized class I TSs responsible for generating farnesane (**1**), bourbonane (**2**), and other various germacrene scaffolds (**3**-**7**) (**Figure 1A**).^9–11^ However, the scaffolds identified thus far are not predicted to be intermediates in the biosynthesis of either laurane or chamigrane-type sesquiterpenoids. Both classes are proposed to arise through the scaffold of bisabolene (**8**), which can undergo further cyclization and modifications to generate the commonly observed in laurane and chamigrane type terpenoids (**9–11**) (**Figure 1B**). To date, no class I TS capable of producing bisabolene or another plausible biosynthetic precursor has been identified in red algae despite exhaustive searches, raising the possibility that an alternative enzyme is responsible for this chemistry.^4,12^ Similar observations in organisms like bacteria and fungi, led to the discovery that enzymes from different families can act as TSs to generate terpene scaffolds.^13,14^ This observation prompted us to question whether red algae employ a non-canonical enzyme to form the key scaffold of bisabolene.

**Figure 1.**
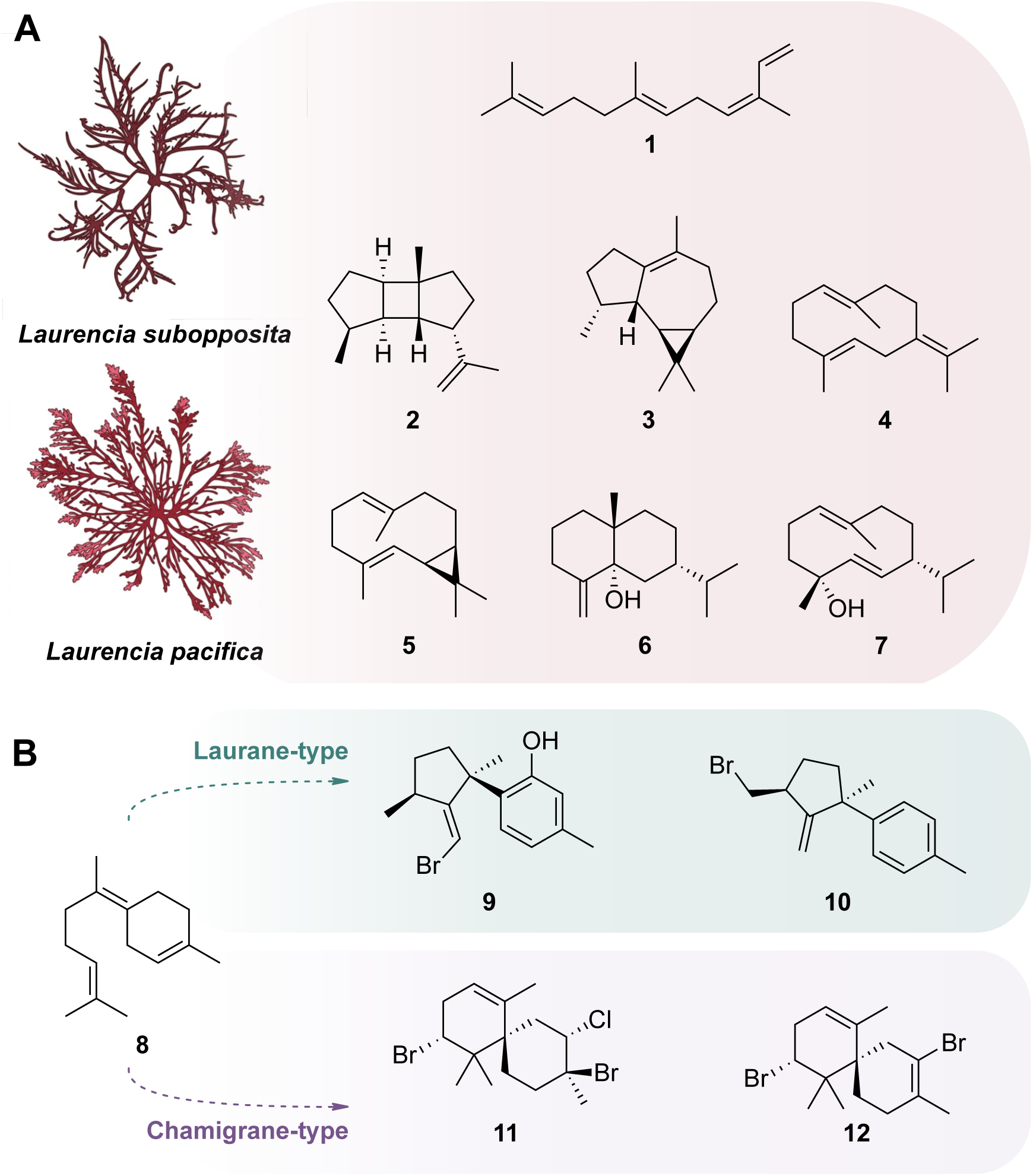
Characterized and proposed products of *Laurencia* terpene biosynthesis. (A) Terpene scaffolds generated by characterized class I terpene synthases from the red algae *Laurencia subopposita* and *Laurencia pacifica*. (B) The bisabolene scaffold (**8**) is proposed to serve as a precursor to representative laurane- and chamigrane-type sesquiterpenoids (**9–12**).

Here, we report the discovery of UbiA-type TSs (UTSs) in red algae that generate terpene scaffolds not previously associated with characterized red algal class I TSs. Most notably, two of the UTSs produce bisabolene, establishing a potential route to both lauranes and chamigrane-type terpenoids. The genes encoding these enzymes are organized within biosynthetic gene clusters (BGCs), further supporting the emerging paradigm that algae and other marine eukaryotes^15–18^ also use proximity-based gene organization for secondary metabolism. Through the functional characterization of UTSs, we establish an alternative route to forming terpene scaffolds previously undescribed in red algae. Together, these findings redefine the enzymatic logic underlying haloterpenoid biosynthesis in red algae and establish UTSs as the core source of the bisabolene family of ecologically important sesquiterpenoids.

## Results

### Genome Mining for UbiA-type Terpene Synthases (UTS)

The cyclization of farnesyl diphosphate (FPP) to bisabolene begins with ionization of the pyrophosphate group through nerolidyl diphosphate (NPP), initiating the carbon-carbon bond formation and ring closure required to generate the bisabolene scaffold (**Figure S1**). Therefore, we reasoned that the enzyme responsible would likely employ chemistry similar to that of class I terpene synthases. Two enzyme families capable of related carbocation-mediated chemistry are isoprenyl diphosphate synthases (IDSs) and UbiA prenyltransferases (PTs). IDSs catalyze the condensation of isopentenyl diphosphate (IPP) and dimethylallyl diphosphate (DMAPP) to generate linear isoprenoid precursors, which then serve as substrates for both PTs and TSs. UbiA-type PTs, on the other hand, are membrane-associated enzymes that prenylate aromatic substrates during the biosynthesis of ubiquinones, menaquinones, and other lipophilic compounds.^19^ Although both enzyme families play prominent roles in primary metabolism, a growing number of examples have shown that these enzymes are capable of forming terpene scaffolds.^20^

Following the formation of bisabolene, the conversion into laurane- or chamigrane-type scaffolds requires an additional cyclization step. We predict that this can occur through the mechanistic capability of class II terpene synthases (TSs), which initiate carbocation formation by protonating an olefinic double bond or epoxide moiety, as observed in the biosynthesis of sesquarterpenoids (C35) derived from tetraprenyl-β-curcumene.^21^ Alternatively, vanadium-dependent haloperoxidases (VHPOs) have shown that they can act analogously to class II TSs, but instead use a halonium ion to achieve ring closure. This has been demonstrated in the biosynthesis of napyradiomycin and merochlorin in bacteria, but also for snyderols from red algae.^22^ Furthermore, several red algal biosynthetic pathways exhibit proximity-based gene organization, including the pathways for kainoid and bromoform biosynthesis and the putative TS and VHPO gene pair identified in *L. subopposita*.^11,17,18^ Taken together, these observations suggested that an alternative bisabolene-forming enzyme could be colocalized with a class II TS or VHPO or another gene involved in terpenoid biosynthesis. Therefore, we searched *Laurencia* genomes for putative genes involved in bisabolene formation, prioritizing candidates located within biosynthetic gene clusters (BGCs).

We used a Hidden Markov Model (HMM)-based approach to search the previously sequenced genomes of *L. subopposita* and *L. pacifica* and to pull out candidate genes from four well-known enzyme families: type II TSs (PF19086), UbiA prenyltransferases (PF01040), VHPOs (PF01569), and polyprenylsynthetases (PF00348).^11^ The results from each search were then queried to determine whether two hits occurred on the same contiguous assembled sequence (contig). All contigs with more than two hits from the different PFAM searches were extracted from the larger genome files and annotated using the Conserved Domain Database (CDD) to identify additional colocalized genes. The resulting annotated sequences were then assigned to putative host or microbial bins based on overall gene cluster architecture and annotation results, and only sequences assigned to the host were retained for further analysis (**Figure S2**).

Across the two species, eight UbiA genes were identified as being clustered with at least one other gene putatively involved in terpenoid biosynthesis, providing candidates for further study (**Figure 2**, **Table S1-S2**). A multiple sequence alignment of all identified UbiA genes revealed two conserved metal-binding motifs, NxxxDxxxD and DxxxD, consistent with motifs observed in canonical UbiA PTs (**Figure S3-4**).^23^ Structure-based similarity searches using AlphaFold-predicted models further supported their assignment as PTs, revealing strong structural conservation with characterized members of the UbiA superfamily (**Figure S5**). The selected UbiA-containing contigs included either isoprenyl diphosphate synthases (IDSs), which could contribute to precursor biosynthesis, or VHPOs or flavin-dependent monooxygenases (FMOs), which could participate in scaffold cyclization or tailoring. Two candidate UbiA genes were identified in *L. pacifica*. LpUTS-1 was colocalized with an IDS, whereas LpUTS-2 was located near an adjacent VHPO. *L. subopposita* contained a greater number of UTS candidates, with six UbiA sequences identified across several contigs, including two nearly identical pairs. The first pair, LsUTS-1a and LsUTS-1b, were both located next to an IDS and a VHPO. The genomic context of LsUTS-2 was distinct, as its contig contained both a VHPO and a nearby FMO. LsUTS-3 encoded a truncated protein due to a premature stop codon and was colocalized with a truncated IDS and a full-length VHPO. Lastly, LsUTS-4a and LsUTS-4b comprised a second nearly identical pair, with each UbiA gene located on a contig containing a neighboring IDS. Phylogenetic analysis of the clusters revealed one set of orthologous genes across both *Laurencia* species (LsUTS-2 and LpUTS-2) (**Figure 2, Figure S6**).

**Figure 2.**
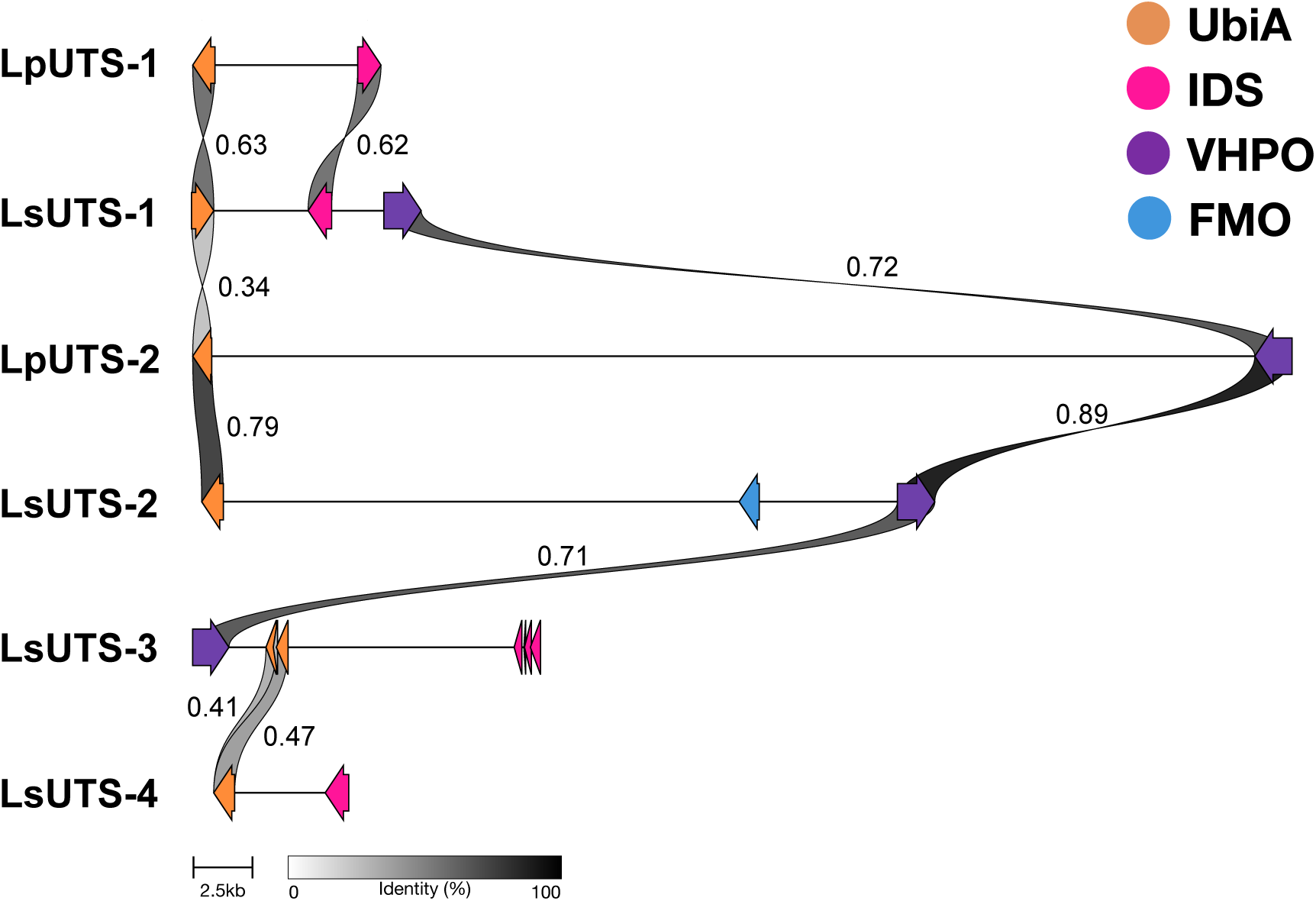
Synteny of putative UTS gene clusters in *Laurencia*. Comparison of genomic regions containing UbiA genes identified in *L. pacifica* (LpUTS) and *L. subopposita* (LsUTS). UbiA genes are shown in orange, isoprenyl diphosphate synthases (IDSs) in pink, vanadium-dependent haloperoxidases (VHPOs) in purple, and flavin-containing monooxygenases (FMOs) in blue. Gray regions connect homologous sequences, with shading corresponding to percent amino acid identity. Values indicate similarity scores between connected regions, and the scale bar represents genomic distance.

### Characterization of UbiA-type TSs using *Nicotiana benthamiana*

UbiA-type enzymes are membrane-associated, which often complicates heterologous expression, particularly in bacterial hosts.^24^ To address this, we utilized the heterologous host *Nicotiana benthamiana*, a system commonly employed for plant biosynthetic pathways to screen activity of the identified gene clusters (**Figure 3A**). Codon-optimized genes from each UbiA-containing gene cluster were individually cloned into pEAQ-HT vectors and transformed into *Agrobacterium tumefaciens.*^25^ Constructs were tested both individually and in combination to assess cluster activity. To enhance terpene precursor availability, all constructs were co-expressed with a construct encoding HMG-CoA reductase (**Figure 3B**).^26,27^

**Figure 3.**
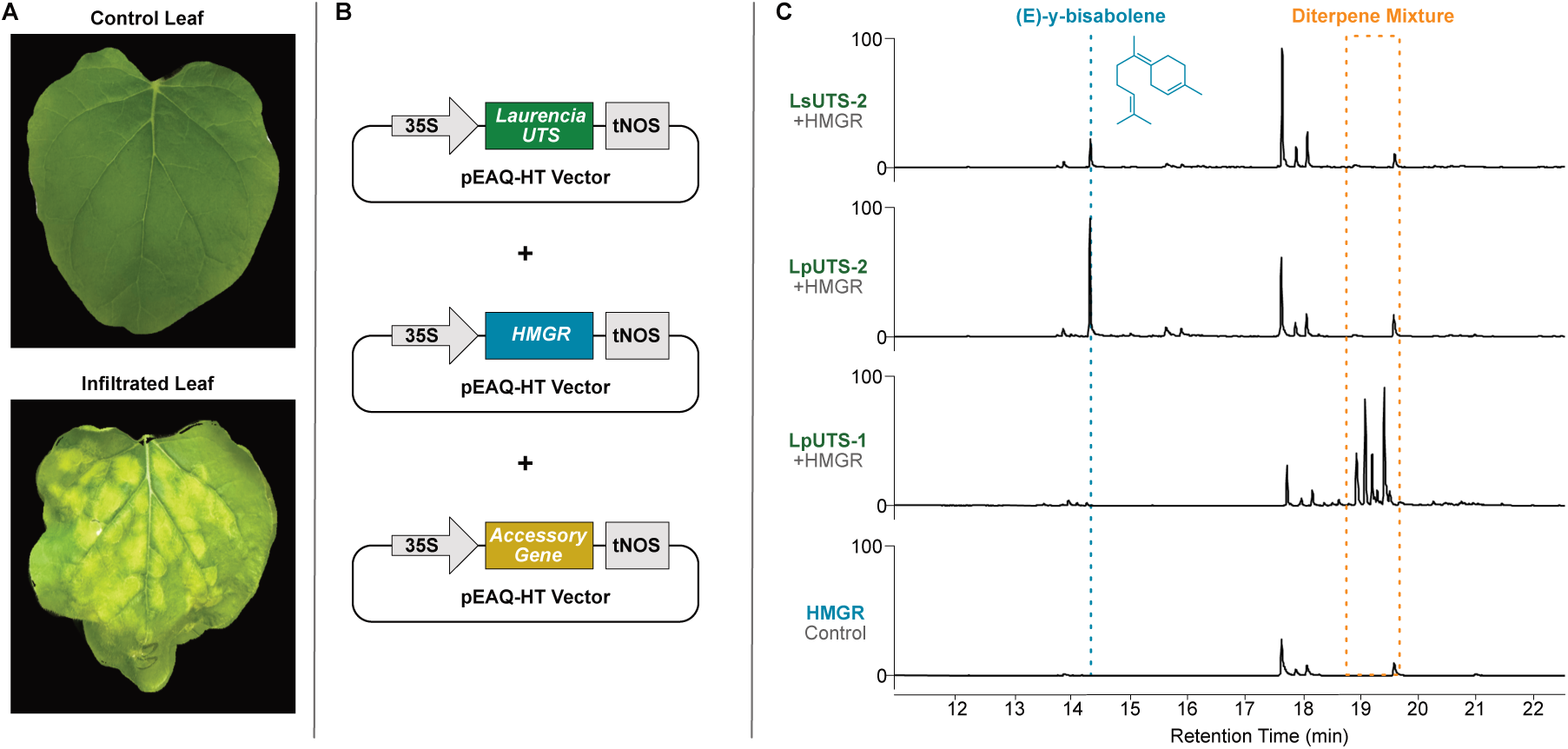
Heterologous expression of red algal UTS enzymes in *Nicotiana benthamiana*. A) Representative *N. benthamiana* leaves comparing an uninfiltrated, control leaf against a leaf infiltrated with agrobacterium containing recombinant protein from *Laurencia* spp. (B) Schematic representation of constructs used in infiltration containing UTS genes along with precursor enhancing HMGR, and accessory genes if present in the cluster. Cauliflower mosaic virus (CaMV) 35S promoter, a CDS, and the nopaline synthase terminator (tNOS) from the pEAQ-HT expression plasmid make up the expression cassette for each tested construct (C) GCMS chromatograms of crude leaf extracts following transient expression.

**Figure 3.**
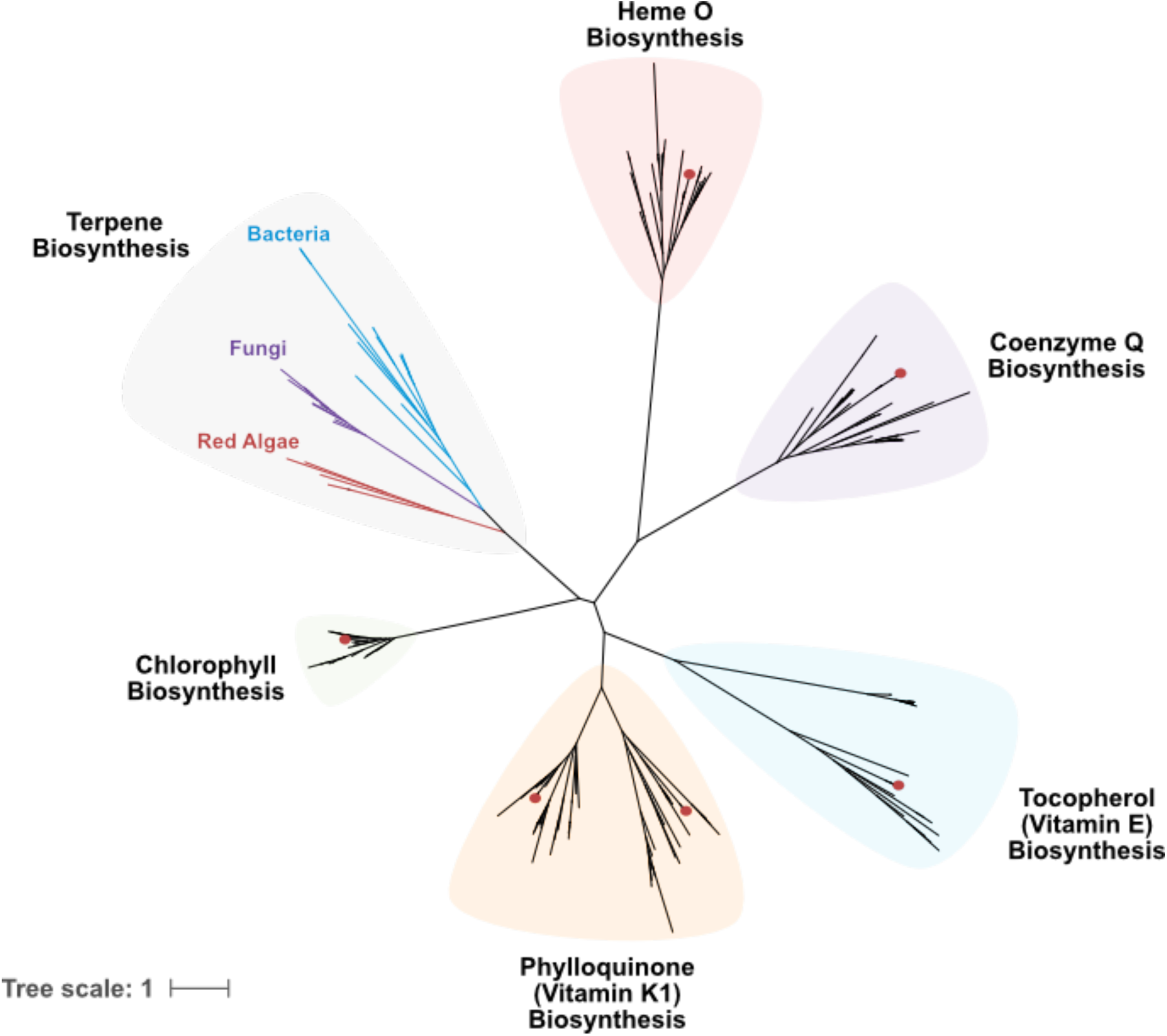
Phylogenetic analysis of UbiA prenyltransferases across Rhodophyta. Maximum-likelihood phylogenetic tree of UbiA genes identified from red algal genomes. The phylogeny resolves five major clades associated with primary metabolic functions, including coenzyme Q, chlorophyll, heme O, vitamin E, and vitamin K1 biosynthesis. A distinct sixth clade contains terpene synthase (TS)-associated UbiAs, including the UbiA-type terpene synthases characterized in this study. Red circles indicate sequences from *L. pacifica* and *L. subopposita* involved in primary metabolism.

The first gene pair, LpUTS-1 and LpIDS-1, produced a mixture of diterpenes (**Fig. 3C, Figure S7**). When tested individually, LpIDS-1 functioned as a geranylgeranyl diphosphate synthase (GGPPS). Titers of diterpene products from LpUTS-1 were improved when coexpressed with LpIDS-1. In *L. subopposita*, two homologous clusters (LsUTS-1a and LsUTS-1b) were identified; however, neither LsUTS-1 nor its associated LsVHPO-1 showed detectable activity in *N. benthamiana*. Despite this, LsIDS-1 was confirmed to also function as a GGPPS (**Figure S8**). Similarly, LsUTS-4a was inactive under these conditions, although the colocalized LsIDS-2 was identified as a functional GGPPS (**Figure S9**).

LsUTS-2 and LpUTS-2 were identified as paralogs across the two *Laurencia* species. LsUTS-2 co-localized with an FMO (LsFMO-1) and a VHPO (LsVHPO-2), while LpUTS-2 paired with LpVHPO-1. When tested individually, both LsUTS-2 and LpUTS-2 produced bisabolene, providing direct evidence for a non-canonical route to this scaffold in red algae (**Figure 2C**, **SI Figure 10-11**). Although no activity was observed for the associated VHPO or FMO enzymes in initial assays, further optimization may be required to capture the activity of these putative tailoring enzymes.

### Sequence and Phylogenetic Analysis of *Laurencia* UbiA-Type Terpene Synthases

Seaweed tissues harbor diverse communities of organisms that engage in various symbiotic interactions with their host (**Figure S12**).^28^ The presence of these symbionts can complicate genome assembly efforts, as DNA from both the host and the symbionts can be incorporated into the resulting assembly. Despite these complications, the resulting assemblies can provide valuable information for assessing gene origin, as interactions between hosts and symbionts can provide opportunities for horizontal gene transfer.^29^ Therefore, to determine the evolutionary origin of the UTSs, we examined the genomic context of the contigs on which they were identified as well as their phylogenetic relationship to primary metabolic UbiAs.

First, we evaluated whether the UTS-containing contigs originated from the *Laurencia* nuclear genomes rather than from microbial contaminants. Multiple features supported their eukaryotic origin, including neighboring genes with homology to eukaryotic proteins, open reading frames separated by relatively large intergenic regions, and assembly coverage, relative abundance, and nucleotide composition consistent with contigs containing established eukaryotic marker genes (**Supplemental File 1**). Because the neighboring IDSs are predicted to supply substrates to the UTSs, we next investigated whether these enzymes were derived from endogenous red algal sequences or were also recruited from outside the red algal lineage. Red algal IDS sequences were retrieved from JGI PhycoCosm and GenBank using the PF00348 profile HMM, together with selected homologs identified in the *L. pacifica* and *L. subopposita* hologenomes. The sequences were aligned with experimentally validated *Arabidopsis thaliana* IDSs, which were included as functional anchor points, and used to construct a maximum-likelihood phylogeny. The number of IDSs per genome identified ranged from 2-6 (**Table S3**) The four IDSs associated with the UTS gene clusters grouped within the red algal GGPPS clade, supporting their origin from within the alga (**Figure S13 and Table S4**). Together, these features provide strong evidence that UTS-containing contigs originate from the red alga host, rather than a microbial contaminant.

Notably, the UTS-containing contigs also contained neighboring sequences associated with mobile genetic elements, including helitron-type transposons. Helitrons are eukaryotic DNA transposons proposed to move through a rolling-circle mechanism and can capture and mobilize neighboring host sequences.^30^ The presence of these mobile elements motivated us to investigate the evolutionary relationships between the UTSs and UbiAs associated with primary metabolism in red algae.

Primary metabolic UbiA sequences were retrieved from Rhodophyta genomes and transcriptomes available through JGI PhycoCosm, NCBI, and GenBank using an HMM based approach (PF01040). Characterized fungal and bacterial UTSs were included as an outgroup. The resulting sequences were aligned and used to construct a maximum-likelihood phylogeny, with clades annotated by BLASTp searches against GenBank non redundant database. The resulting phylogeny showed six major clades including a UTS associated clade and five clades that were assigned to the biosynthesis of primary metabolites: HemeO, Coenzyme Q, Chlorophyll, Vitamin E, and Vitamin K1 biosynthesis (**Figure 4, Figure S14, and Table S5**). Across the red algal genomes examined, at least five UbiA genes corresponding to primary metabolic functions were consistently identified, showing that non-terpenoid-producing species contained an average of five copies per genome (**Table S6**). In addition to primary metabolic UbiAs, the *Laurencia* genomes contained additional UbiA genes corresponding to the UTSs characterized here. These specialized UbiAs form a monophyletic clade with both bacterial and fungal UTSs and show no close phylogenetic relationship to the primary-metabolic red algal UbiAs. Alongside the presence of mobile genetic elements on the contigs, these findings support that the *Laurencia* UTSs originated from outside of the red algal lineage. The fact that these UTSs are only found in *Laurencia* spp. and not any other red algal genera whose sequencing data were available corresponds well to the fact that halogenated laurane and chamigrane terpenoids are specific to *Laurencia* spp. within red algae.^3^

Taken together, these phylogenetic analyses suggest that the UTS-containing gene clusters have a mixed evolutionary origin, with the TS component likely acquired from outside the red algal lineage while the associated IDS genes appear to be endogenous to red algae.

## Discussion

Here we report the first identification of red algal UTSs from the genus *Laurencia*, previously only observed within fungi, bacteria, and most recently sponges.^20,31^ The discovery of these non-canonical enzymes was made possible by their presence in biosynthetic gene clusters, a feature increasingly recognized in eukaryotic genomes.^15,16,32^ By prioritizing gene candidates that were located on contigs containing one or more additional putative biosynthetic genes, we identified eight gene clusters across *L. pacifica* and *L. subopposita.* Using the heterologous host, *N. benthamiana*, we linked these gene clusters as being involved in terpenoid biosynthesis by demonstrating the formation of terpene scaffolds. The characterization of two UTSs that produce bisabolene is notable because this scaffold was reported to be missing among the previously characterized canonical class I TSs, signifying red alga has evolved at least two convergent routes to producing scaffolds for haloterpenoids.^11^

The bisabolene scaffold is not unique to red algae, but instead is widely observed across diverse lineages. Plants, bacteria, and fungi have all been shown to produce various isomers, either α-, β-, and γ-bisabolene. While all these isomers share a C1–C6 bond that forms the ring, they differ in where the deprotonation occurs. Furthermore, the enzymes that produce them also appear to vary across lineages. In plants, several class I TSs have been characterized including synthases that produce the following isomers: (E)-α-bisabolene, (S)-β-bisabolene, and (Z)-γ-bisabolene. In grand fir, (E)-α-bisabolene synthase supplies a precursor to todomatuic acid and juvabione, compounds associated with defense.^33^ The actinomycete *Cryptosporangium arvum* DSM 44712 also encodes a canonical class I TS that produces (Z)-γ-bisabolene.^34^ In contrast, insects have been shown to use IDS-like TSs to produce bisabolenes. The stink bug *Nezara viridula*, uses an IDS-like TS to produce (+)-(S,Z)-α-bisabolene, which is subsequently modified into sex pheromones in male insects.^35^ Fungi provide a closer enzymatic parallel to red algae because they have several UTSs. In *Antrodia cinnamomea*, a UTS, Tps1a, produces (+)-(S,Z)-α-bisabolene and appears in a cluster containing tailoring genes.^36^ As in our analysis of *Laurencia* genomes, the UTSs were discovered only after the class I TS candidates had been exhausted. The (E)-γ-bisabolene-producing UTS characterized here extends the known product range of this enzyme family. The production of bisabolene isomers by class I TSs, UTSs, and IDS-related enzymes shows that this comparatively simple monocyclic scaffold is accessible through distinct protein architectures and could represent an early evolved product in terpene biosynthesis. The integral membrane nature of the UTSs highlights subcellular localization as a potentially important factor in *Laurencia* terpenoid biosynthesis, with implications for substrate access and interactions with downstream tailoring enzymes. Determining their native localization could help clarify whether terpenoid biosynthesis is spatially coupled to the vesicular transport and extracellular release of halogenated metabolites reported in *Laurencia*.^37^

The *Laurencia* UTSs also raise questions about the origin of red algal terpene pathways, particularly because they occur alongside canonical class I TSs. The presence of both routes suggests red algae have maintained multiple ways to produce terpene scaffolds. In our phylogenetic analysis, the red algal UTSs formed a distinct group from canonical UbiA enzymes and clustered with terpenoid biosynthetic enzymes from fungi and bacteria. This pattern is consistent with an origin outside the red algal lineage, although the phylogeny alone cannot establish how these genes were acquired. Retroviral elements on contigs containing the UTS gene clusters raise the possibility that mobile genetic elements played a role in their evolution, but their proximity does not demonstrate a role in gene transfer.

The absence of characterized enzymes producing the chamigrane and laurane scaffolds, raises the possibility of a biosynthetic route for both classes from a bisabolene intermediate through the activity of additional tailoring enzymes.^12^ So far, no wild-type enzyme has been identified to produce the laurane or chamigrane scaffolds, though mutant versions of the class I TS, SydA, have been shown to produce chamigranes.^38^ Therefore, we hypothesize that this must occur through tailoring enzymes in red algae. Cyclization through tailoring enzymes has previously been observed in *Laurencia obtusa*, in which a vanadium-dependent bromoperoxidase (V-BrPO) catalyzed the bromination and cyclization of nerolidol to form a mixture of constitutional isomers of snyderol.^39,40^ This precedent suggests that the co-localized vanadium-dependent haloperoxidases (VHPOs) could facilitate a similar scaffold transformation, providing a possible route from bisabolene to the more complex cyclic products. The VHPO genes located near some of the UTSs are therefore candidates for downstream transformations. Whether they accept the bisabolene products reported here, and whether those reactions lead toward laurane or chamigrane metabolites, remains to be tested.

Despite their independent evolutionary origin, class I TSs and UTSs share a conserved catalytic logic and ultimately both generate terpene scaffolds despite arising independently through evolutionarily distinct protein families. The emergence of parallel biosynthetic strategies may reflect strong selective pressure for terpene production, with each pathway contributing distinct scaffolds that fulfill specialized biological functions. Discovery of these non-canonical pathways, together with emerging evidence for biosynthetic gene clustering in red algae, represents a paradigm shift in our understanding of eukaryotic secondary metabolism and provides a roadmap for future genome-mining efforts to uncover the full extent of terpene biosynthetic diversity

## Acknowledgments

We thank UC San Diego colleagues L. Aluwihare for access to GCMS instrumentation and A. Mrse for access to NMR equipment. The authors also acknowledge A. Papikian and J. Pacheco (Salk Institute) for assistance in growing *N. benthamiana* in early feasibility studies. This research was supported by NIH grant R01-GM146224 to B.S.M., the Tribal Membership Initiative Fellowship (UC San Diego) to H.K.B., the Scripps Institutional Postdoctoral Fellowship to A.J.B., the NSF Graduate Research Fellowship Program to T.S.S., and the Department of Energy Joint Genome Institute (CSP-504335 to B.S.M.).

## Supporting Information

### Materials and Methods

#### General Methods

##### GCMS Analysis

GCMS analysis was performed on an Agilent 7890A gas chromatograph with an Agilent 5975C mass spectrometer using an Rtx-5Sil 30 m × 0.25 mm, 0.25-µm column. The oven program was as follows: hold 70 °C for 3 min, 10 °C min^−1^ to 325 °C and hold at 325 °C for 3 min. The flow rate was 1 mL min^−1^, and the injection volume was 1 µl. The splitless injection inlet temperature was 270 °C, the MS transfer line temperature was 280 °C, the MS source temperature was 230 °C and the MS quad temperature was 150 °C. Data were analyzed using Masshunter.

#### Methods

##### Phylogeny of UbiA Gene Families

Ubi-A family genes were identified by performing HMM searches with pattern PF01040 against predicted proteins from previously described metagenomic assemblies of *Laurencia pacifica* and *Laurencia subopposita*^1^ as well as Rhodophyta reference genomes downloaded from JGI Phycocosm (https://phycocosm.jgi.doe.gov/phycocosm/home/releases?flt=rhodophyta) and the NCBI Genomes database. A more extensive search for sequenced individual *Laurencia* Ubi-A protein relatives from all taxonomic groups was performed by BLASTP search against GenBank nr (accessed November 2025). Combined UbiA family proteins from all sources were dereplicated at (100%?) identity using CD-Hit v4.8.1,^2^ put into multiple sequence alignments with MUSCLE v3.8 (Edgar 2004),^3^ then used to construct maximum likelihood phylogenetic trees using RaxML v1.2.2,^4^ with model JTT+G, rooted using the fungal dimethylallyltryptophan synthase (DMATS) gene from *Claviceps purpurea*. Database sources for the sequences in this tree are described in Table S5.

##### Eukaryotic Contig Identification

Because *Laurencia* MAGs were recovered from metagenomic enrichments rather than axenic cultures, the taxonomic origins of genomic contigs containing Ubi-A sequences were assessed using multiple independent criteria. Relative assembly abundances in metagenomic DNA samples were estimated by calculating contig coverages using assembly read recruitment with BOWTIE2 v2.5.4,^5^ and the idxstats function of SAMTOOLS v1.22.^6^ Gene spacing distances and neighborhood context of contigs containing targeted Ubi-A sequences were determined using manually curated annotations of all potential open reading frames on the same contig, as identified by Prokka 1.14.6^7^ and MetaEuk v6 predictions of gene locations.^8^ Functional domain activities within each predicted protein were assigned based on manual curation of results from InterProScan v5.77-108.0^9^ and closest BLASTP matches in the GenBank nr database.

##### Metatranscriptomic Assembly

Metatranscriptomic assemblies were performed separately for each *Laurencia* sample using SPADES v4.20^10^ from Illumina reads deposited in the GenBank SRA database under accession numbers SRR25727165 (*L. subopposita*) and SRR25726911 (*L. pacifica*). Reads were trimmed and quality filtered with Trimmomatic-0.39 (Bolger 2014) using the following parameters: ILLUMINACLIP, TruSeq3-PE.fa:2:30:10: HEADCROP:11, SLIDINGWINDOW:4:20, MINLEN:130. Prior to assembly, reads from *L. subopposita* were down-sampled to a total of 100 million pairs, approximately matching the number available from *L. pacifica.* Transcriptome assembly completeness was assessed using BUSCO v6.0.0 ^11^ with lineage reference rhodophyta_odb12. Assembled contigs were mapped to UbiA protein sequences using BLASTX searches. Relative transcriptomic abundances of UbiA genes were assessed by mapping unassembled transcriptome reads to assembled contigs using BOWTIE2 v2.5.4 and the idxstats function of SAMTOOLS v1.22.

##### Plant Growth Conditions

*N. benthamiana* were grown in a Percival Biological Incubator (Model I-36LL) at 25 C under 16-h light/8-h dark cycles and watered once a week with distilled water. Experiments were conducted on 5-week-old plants.

##### Transient Expression of Algal Genes in *N. benthamiana*

Cultures containing genes of interest were individually transformed into electrocompetent Agrobacterium cells (Strain GV3101) and grown for 72h. Single colonies were used to inoculate a 30mL culture that then grew for 48h. On morning of transformation, cells were centrifuged at 5,000g for 5 min, then the supernatant was discarded and resuspended to an OD600 =2.0 in the infiltration buffer (10 mM MES pH 5.6, 10 mM MgCl_2_ and 150 μM acetosyringone) and incubated for two hours at room temperature. Genes in cultures were tested by mixing individual cultures together. Infiltration occurred on the underside of *N. benthamiana* leaves on different plants (n=3) and were collected 5 days after infiltration. For volatile sesquiterpenes, plants were removed from incubator after infiltration and left on bench at room temperature to avoid loss by high humidity in growth chamber.

##### Extraction from *N. benthamiana*

The entire leaf that was infiltrated was frozen in liquid nitrogen and homogenized using a mortar and pestle. Ground tissue was extracted with 2mL of 2:1 methanol/dichloromethane and stirred overnight. Samples were then resuspended with 8mL of distilled water and extracted with 2mL of hexane. Hexane fractions were filtered and subjected to GC-MS analysis.

### Supplementary Tables

**Table S1:**
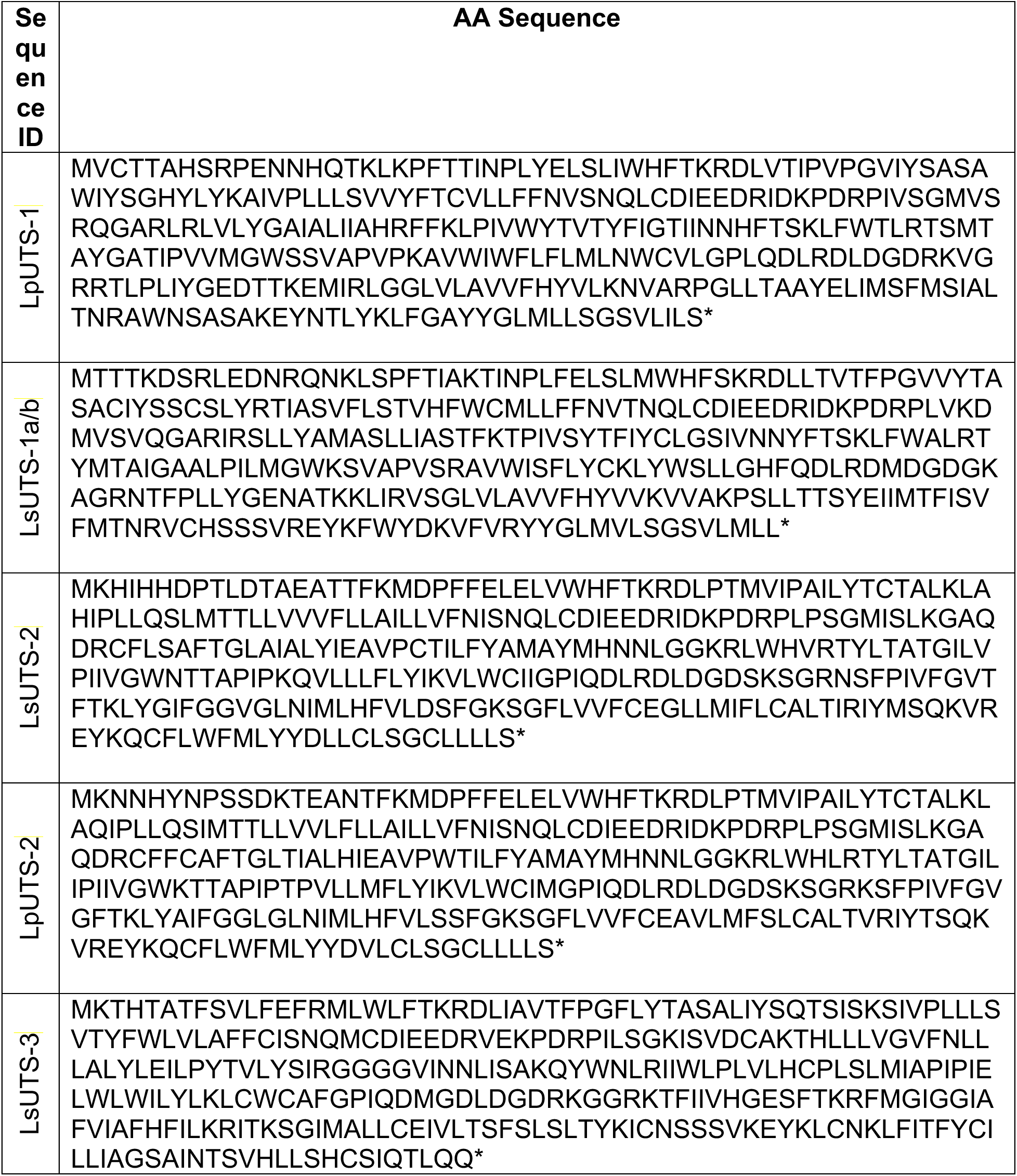
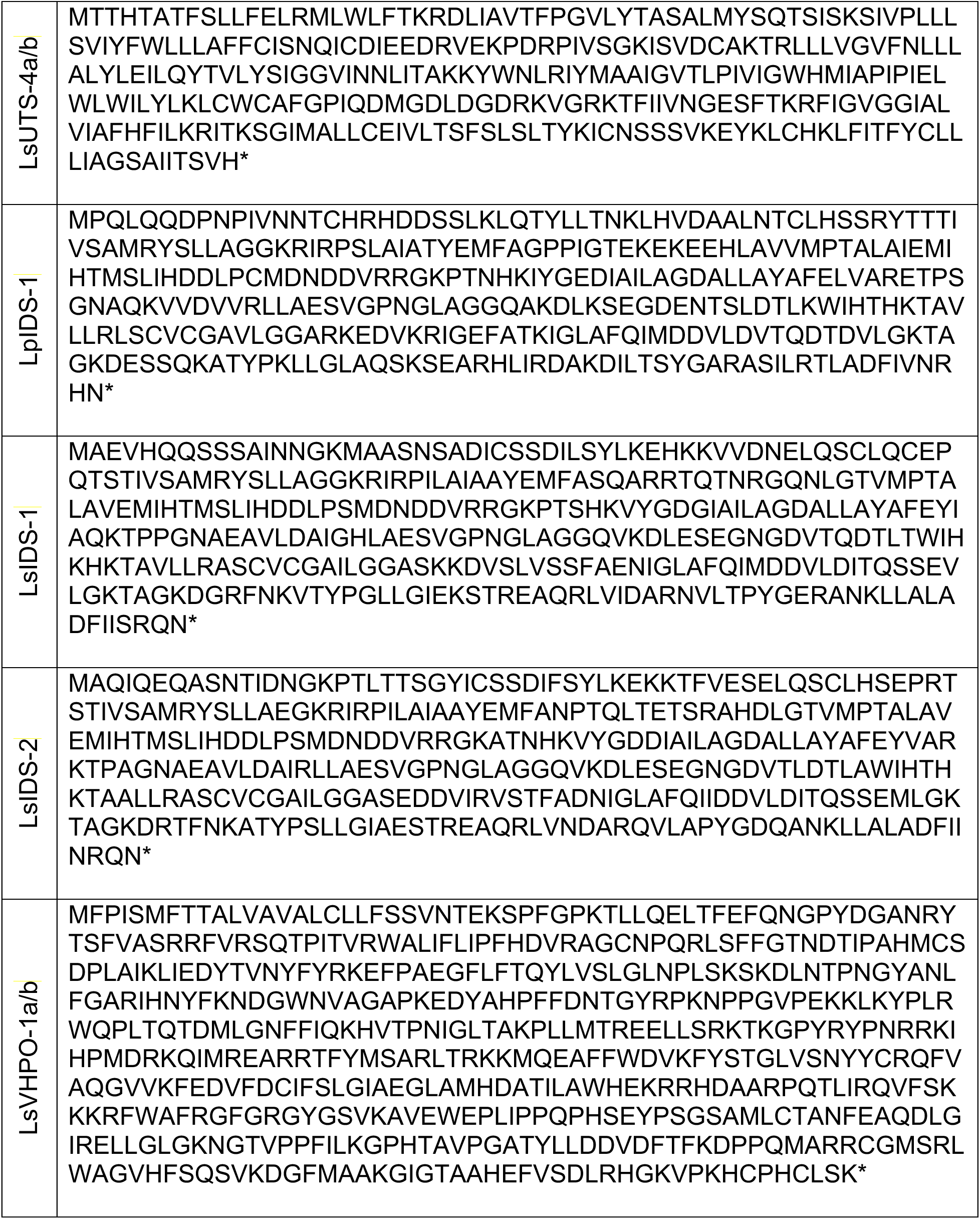
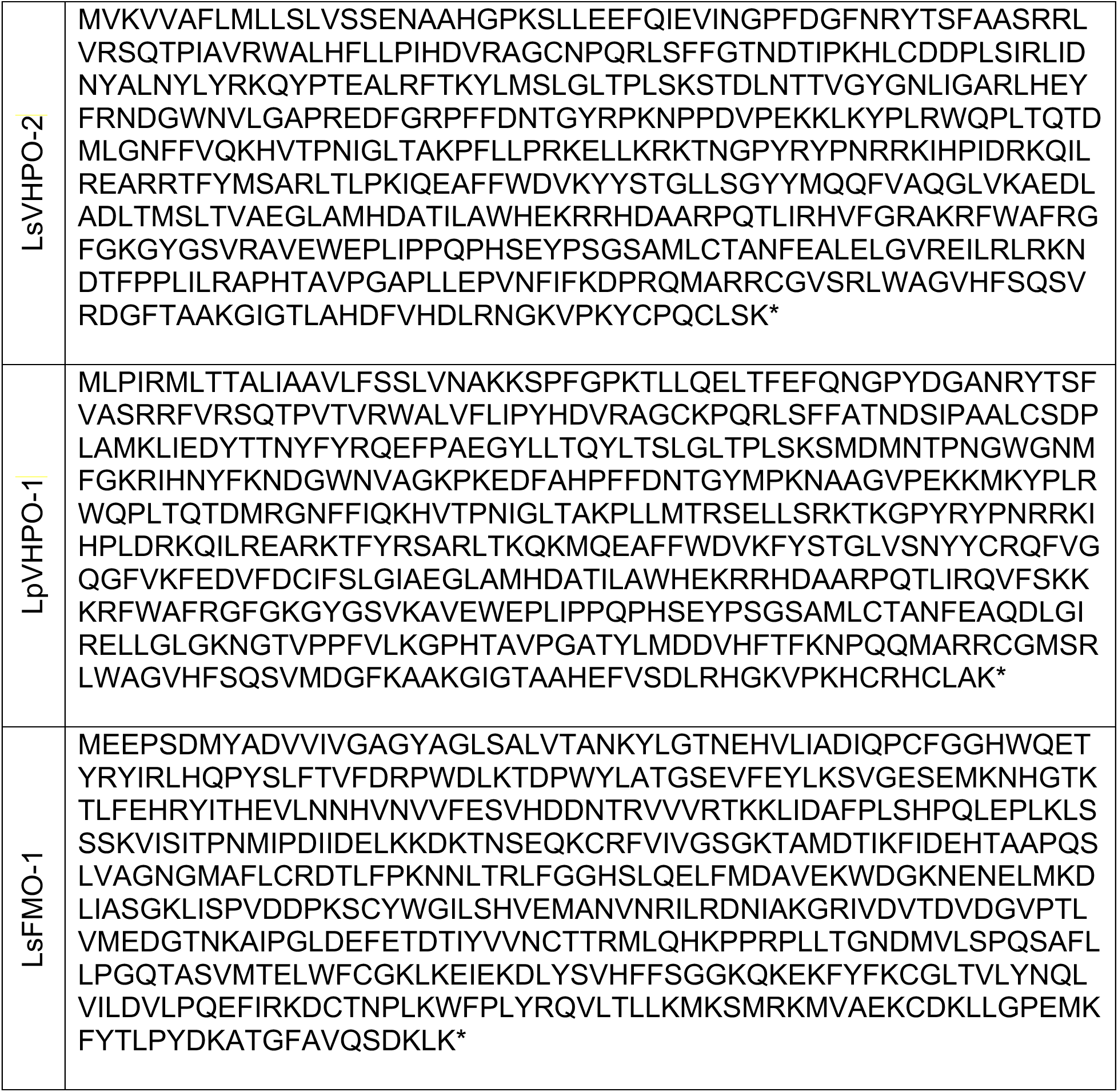
Gene Sequences of Identified UTS Gene Clusters.

**Table S2.** Identified UbiA genes and their surrounding neighbors with predicted functions.

| **UbiA Sequence** | **Accompanying Gene(s)** | **Predicted Function(s)** |
| --- | --- | --- |
| LpUTS-1 | LpIDS-1 | Isoprenyl diphosphate synthase |
| LsUTS-1a | LsIDS-1a,<br>LsVHPO-1a | Isoprenyl diphosphate synthase, vanadium-dependent haloperoxidase |
| LsUTS-1b | LsIDS-1b,<br>LsVHPO-1b | Isoprenyl diphosphate synthase, vanadium-dependent haloperoxidase |
| LpUTS-2 | LpVHPO-1 | vanadium-dependent haloperoxidase |
| LsUTS-2 | LsFMO-1,<br>LsVHPO-2 | Flavin-dependent monooxygenase, vanadium-dependent haloperoxidase |
| LsUTS-3 | LsIDS-2,<br>LsVHPO-3 | Isoprenyl diphosphate synthase, vanadium-dependent haloperoxidase |
| LsUTS-4 | LsIDS-3a | Isoprenyl diphosphate synthase |
| LsUTS-4b | LsIDS-3b | Isoprenyl diphosphate synthase |

**Table S3.** Number of IDS Genes in Queried Red Algal Genomes.

| Red algal species | num<br>PF00348<br>matches |
| --- | --- |
| *Cyanidioschyzon merolae strain 10D* | 6 |
| *Rhodomonas marina* | 4 |
| *Galdieria sulphuraria MtSh* | 4 |
| *Galdieria sulphuraria 002* | 4 |
| *Chondrus crispus Stackhouse* | 4 |
| *Porphyridium purpureum CCMP1328* | 3 |
| *Gracilariopsis chorda SKKU-2015* | 3 |
| *Gracilaria vermiculophylla HapMaleFtJ-2017* | 3 |
| *Gracilaria gracilis GNS1m* | 3 |
| *Gracilaria chilensis NLEC103-M9* | 3 |
| *Gracilaria caudata M-176 S67* | 3 |
| *Galdieria yellowstonensis 108.79 E11* | 3 |
| *Galdieria sulphuraria RT22* | 3 |
| *Galdieria sulphuraria MS1* | 3 |
| *Galdieria sulphuraria Azora* | 3 |
| *Galdieria sulphuraria 5572* | 3 |
| *Galdieria sulphuraria 074W* | 3 |
| *Galdieria phlegrea Soos* | 3 |
| *Cyanidiococcus yangmingshanensis 8.1.23 F7* | 3 |
| *Porphyra umbilicalis isolate 4086291* | 2 |
| *Laurencia subopposita* | 2 |
| *Gracilaria domingensis SP 400837* | 2 |
| *Galdieria sulphuraria YNP5578.1* | 2 |
| *Cyanidium caldarium DBV 063 E5* | 2 |

**Table S4.**
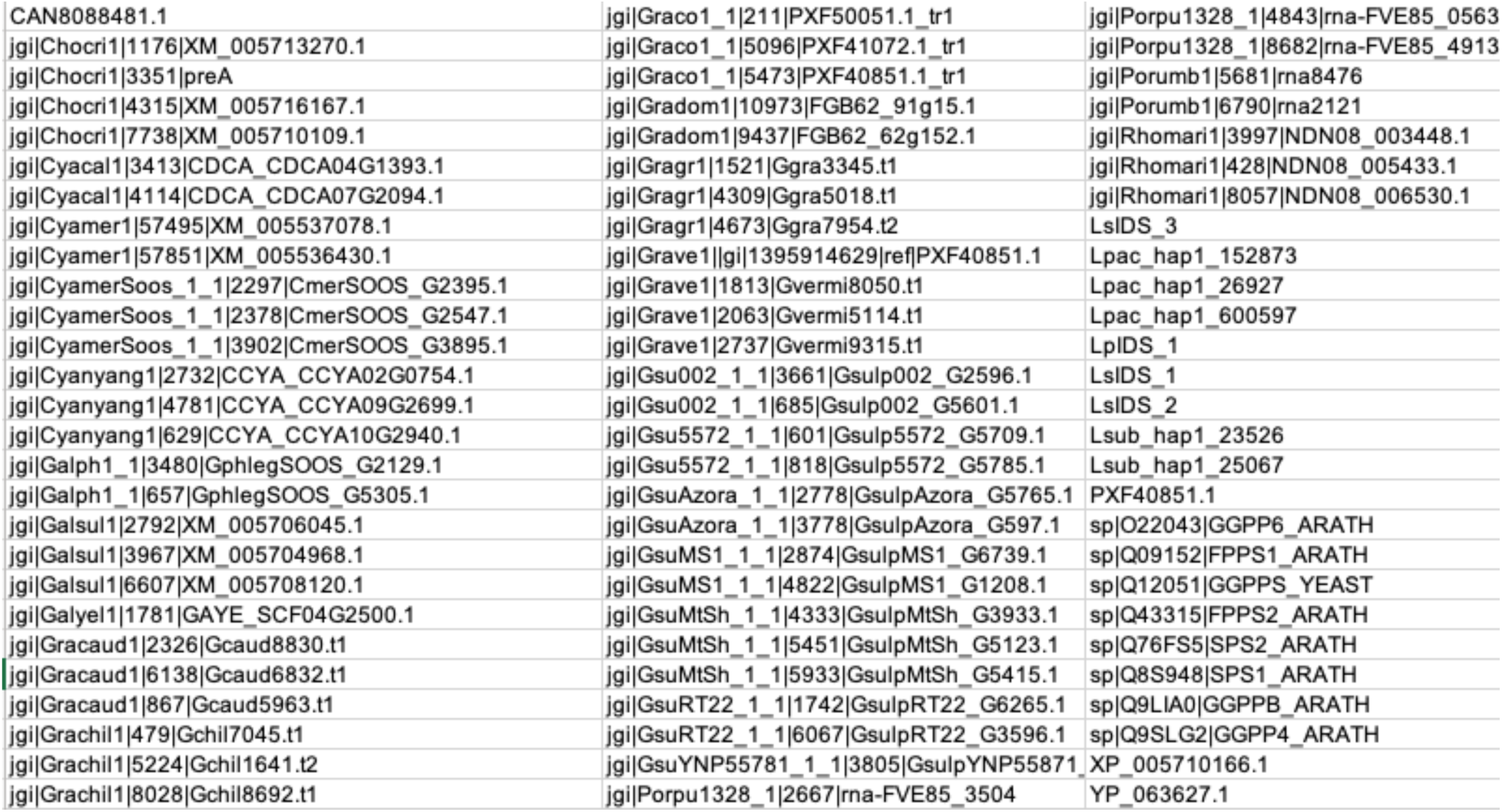
Sequences used in IDS Phylogenetic Tree.

**Table S5.**
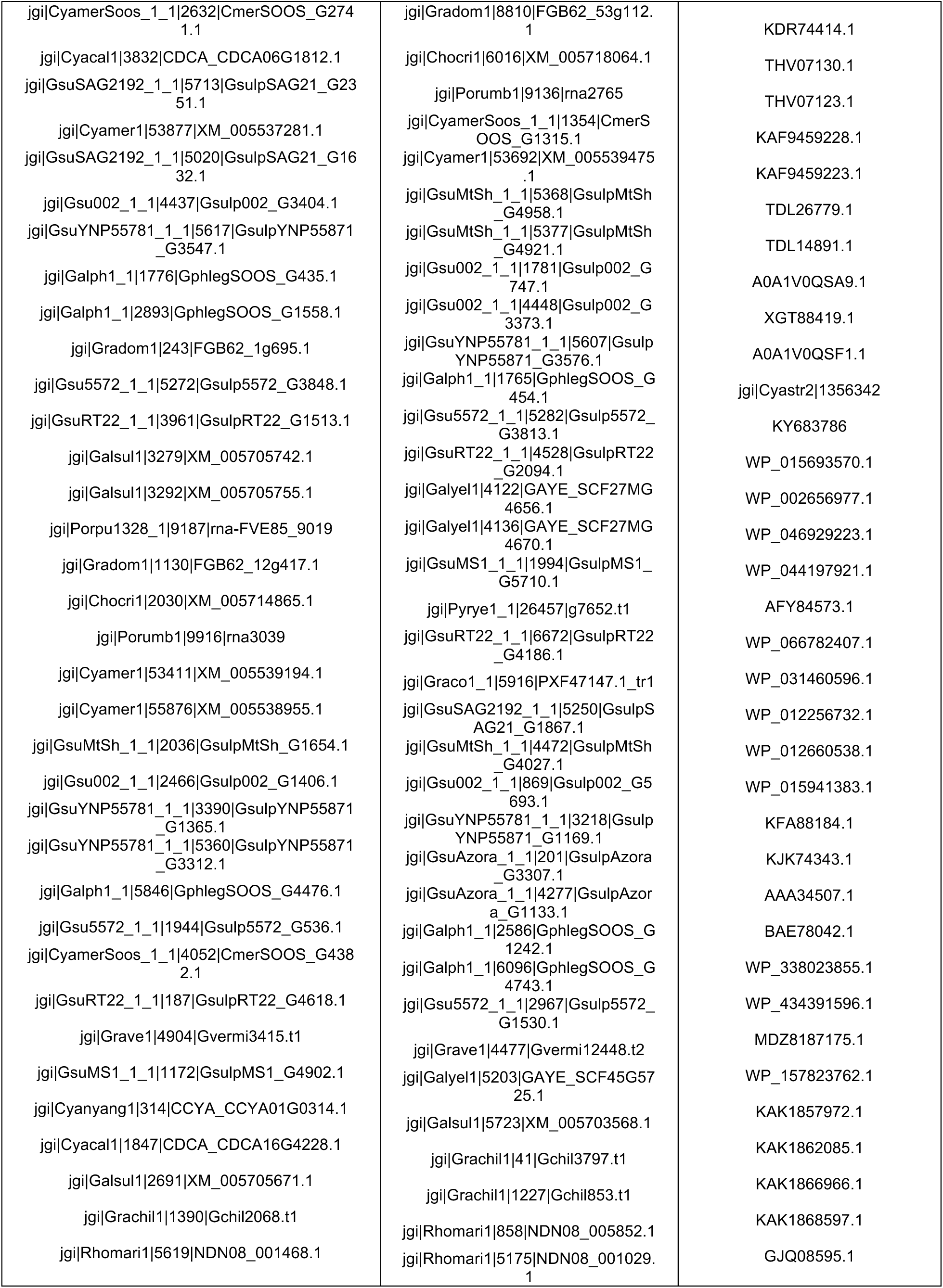
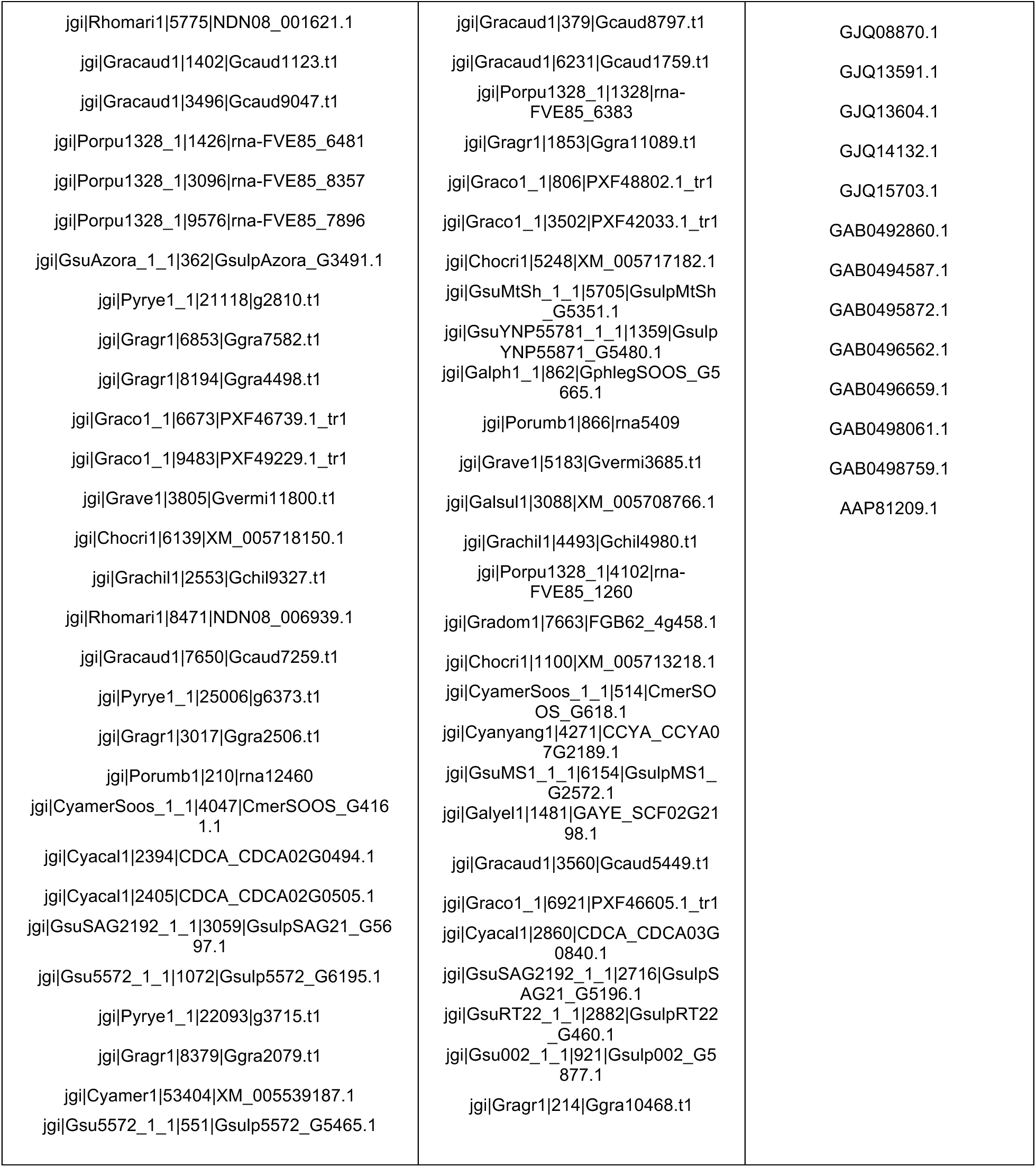
Sequences used in UbiA Phylogenetic Tree.

**Table S6.** Number of Primary Metabolic UbiA Genes in Queried Red Algal Genomes.

| Species name | PF01040 (UbiA prenyl transferase) |
| --- | --- |
| *Chondrus crispus* Stackhouse | 5 |
| *Cyanidium caldarium* DBV 063 E5 | 5 |
| *Cyanidioschyzon merolae* Soos | 5 |
| *Cyanidioschyzon merolae* strain 10D | 5 |
| *Cyanidiococcus yangmingshanensis* 8.1.23 F7 | 5 |
| *Galdieria phlegrea* Soos | 7 |
| *Galdieria sulphuraria* 002 | 6 |
| *Galdieria yellowstonensis* 108.79 E11 | 6 |
| *Gracilaria caudata* M-176 S67 | 6 |
| *Gracilaria chilensis* NLEC103-M9 | 6 |
| *Gracilariopsis chorda* SKKU-2015 | 7 |
| *Gracilaria domingensis* SP 400837 | 4 |
| *Gracilaria gracilis* GNS1m | 7 |
| *Gracilaria vermiculophylla* HapMaleFtJ-2017 | 4 |
| *Galdieria sulphuraria* 074W | 6 |
| *Galdieria sulphuraria* 5572 | 6 |
| *Galdieria sulphuraria* Azora | 6 |
| *Galdieria sulphuraria* MS1 | 5 |
| *Galdieria sulphuraria* MtSh | 5 |
| *Galdieria sulphuraria* RT22 | 7 |
| *Galdieria sulphuraria* SAG21.92 | 5 |
| *Galdieria sulphuraria* YNP5578.1 | 6 |
| *Porphyridium purpureum* CCMP1328 | 6 |
| *Porphyra umbilicalis* isolate 4086291 | 4 |
| *Neopyropia yezoensis* U-51 | 4 |
| *Rhodosorus marinus* CCMP1338 | 6 |
| *Pyropia yezoensis* | 5 |
| *Galdieria partita* | 6 |
| *Galdieria yellowstonensis* | 6 |
| *Pyropia vietnamensis* | 8 |

### Supplementary Figures

**Figure S1.**
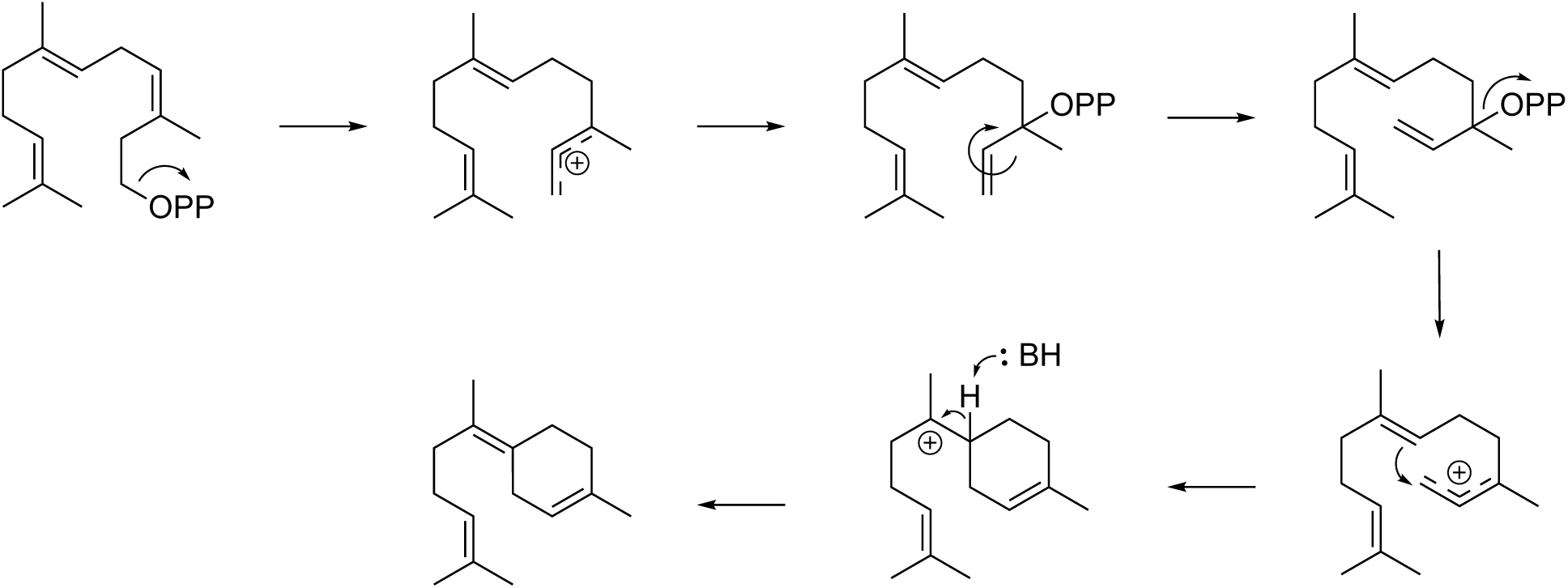
Bisabolene Biosynthesis. Biosynthetic scheme from farnesylpyrophosphate (FPP) through nerolidyl diphosphate to γ-bisabolene.^12,13^

**Figure S2.**
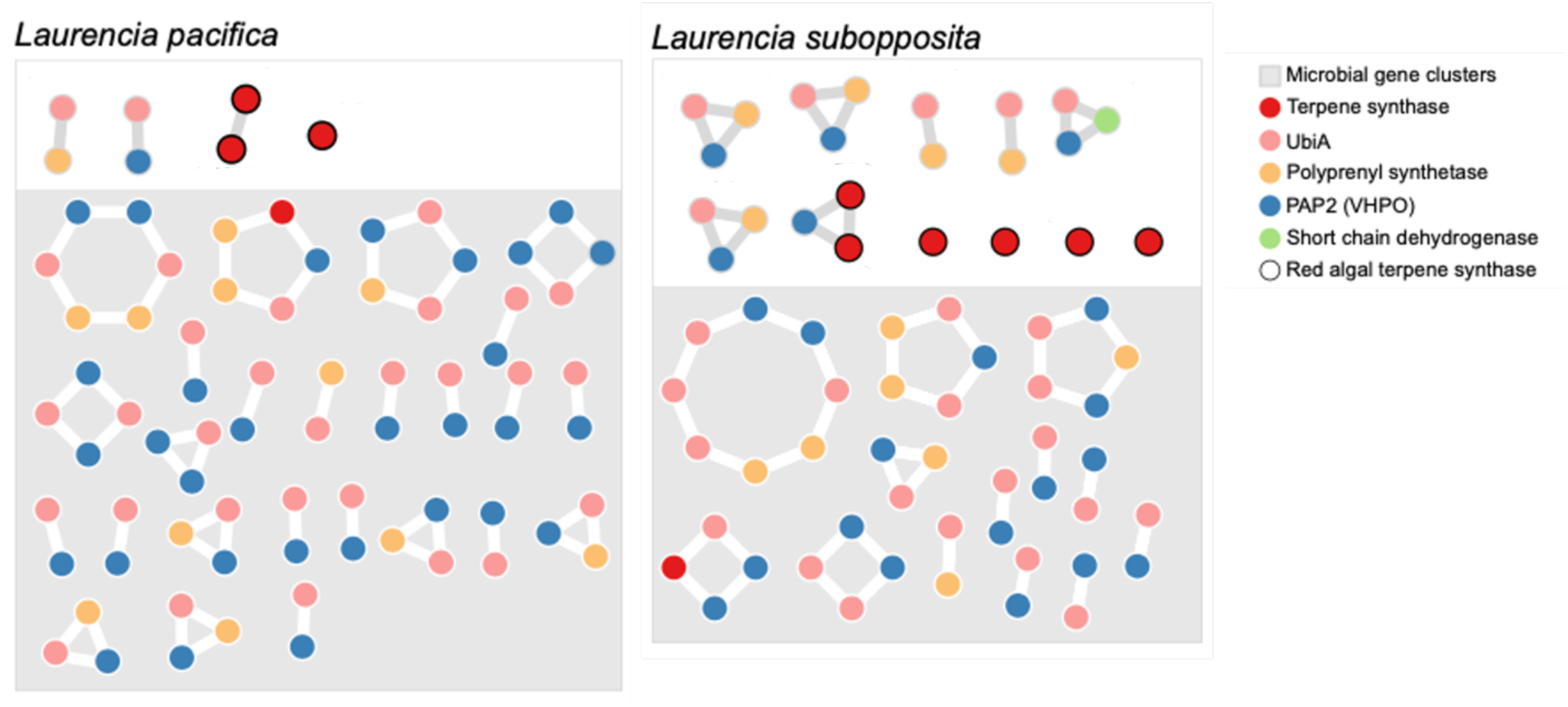
Gene co-occurrence network of *Laurencia* genes. Each dot represents a gene, and each connection indicates physical co-occurrence on a contiguous sequence without respect to order or distance. Contigs from microbial sources are in grey boxes.

**Figure S3.**
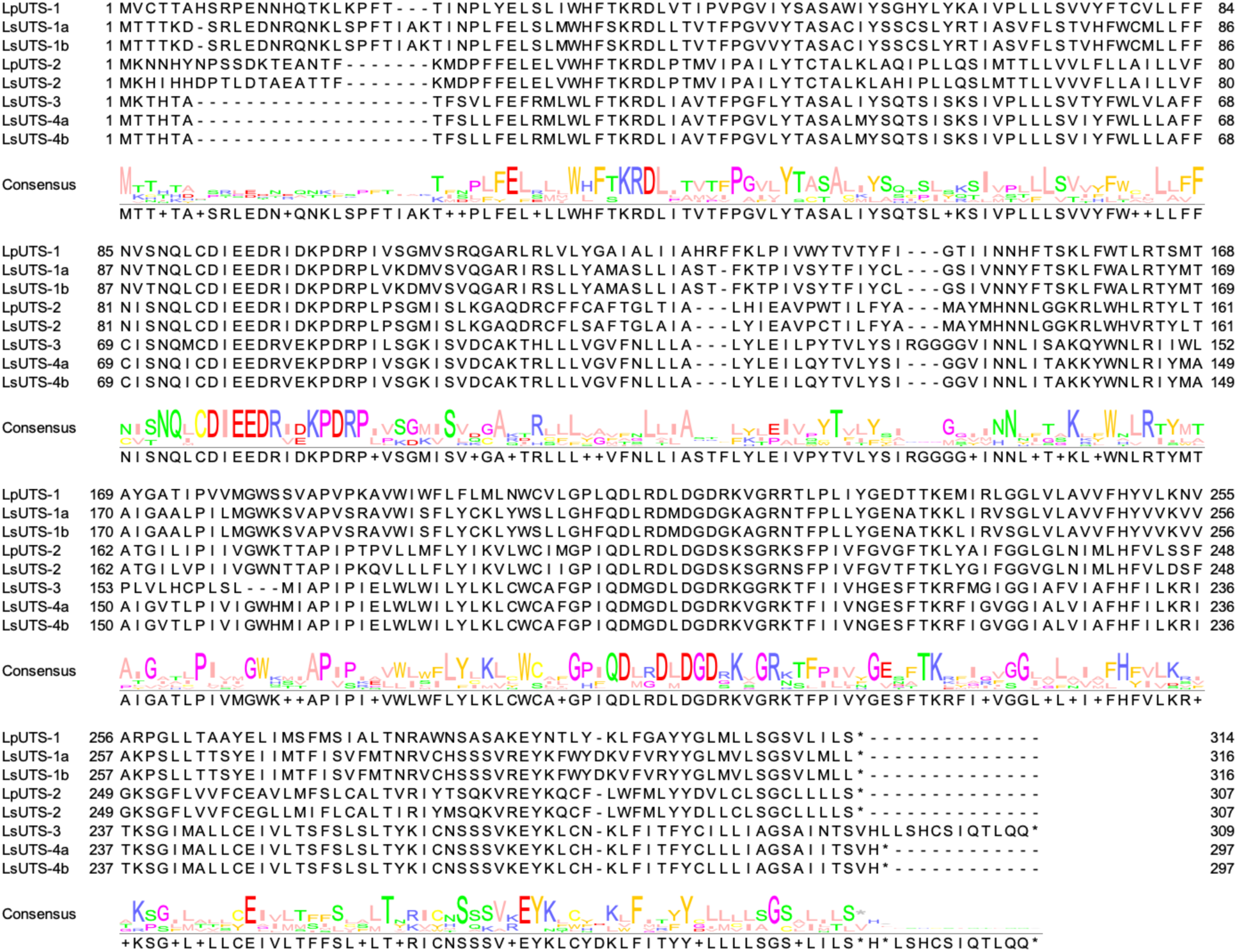
Multiple sequence alignment of *Laurencia* UTSs. Amino acid sequences of UbiA homologs identified in *L. pacifica* and *L. subopposita* were aligned to assess sequence conservation across the family. Residues are colored according to amino acid properties, and sequence conservation at each position is represented by the consensus logo below the alignment. Logo made using Jalview software.^14^

**Figure S4.**
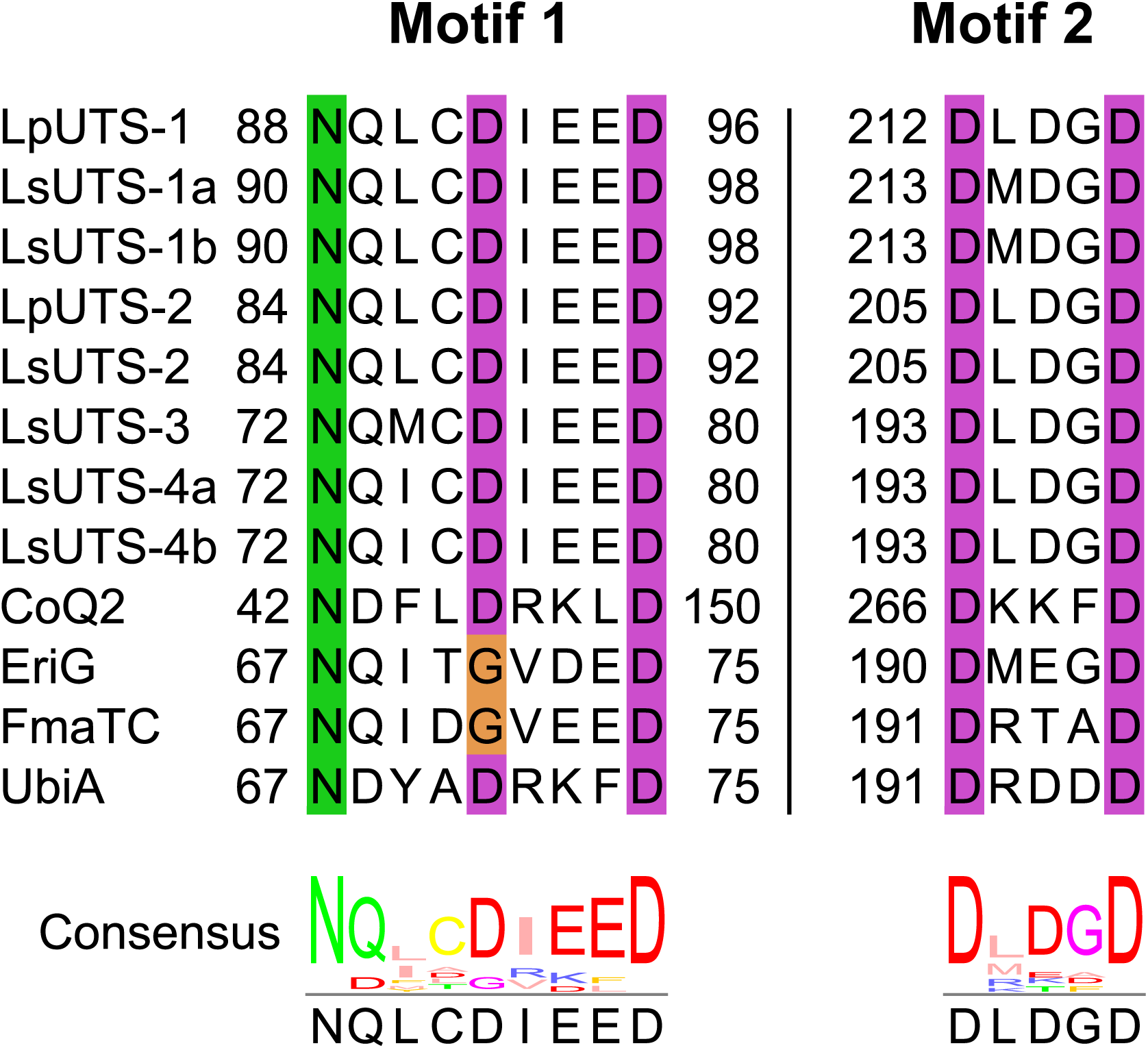
Conservation of catalytic motifs in *Laurencia* UbiA proteins. Alignment of the two conserved aspartate-rich motifs in the identified *Laurencia* UbiA sequences with those of characterized UbiA-family prenyltransferases and UTSs. The *Laurencia* proteins retain the conserved NxxxDxxxD and DxxxD motifs associated with UbiA-family catalysis. Residue numbers indicate the positions of each motif within the corresponding protein sequence.

**Figure S5.**
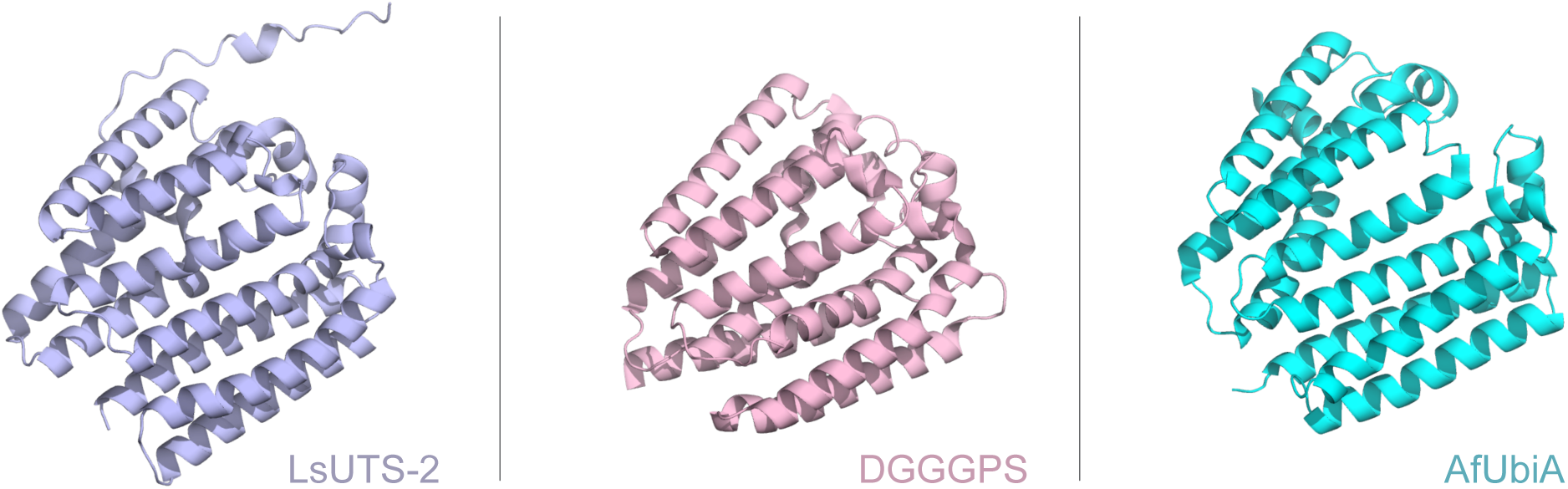
Structural model of LsUTS-2 and its closest structural homologs. The predicted structure of LsUTS-2 was queried against the Protein Data Bank using DALI^15^. DGGGPS (PDB: 6M31) and AfUbiA (PDB: 4TQ4), shown here as ribbon representations, were the two closest structural matches identified by the search.

**Figure S6.**
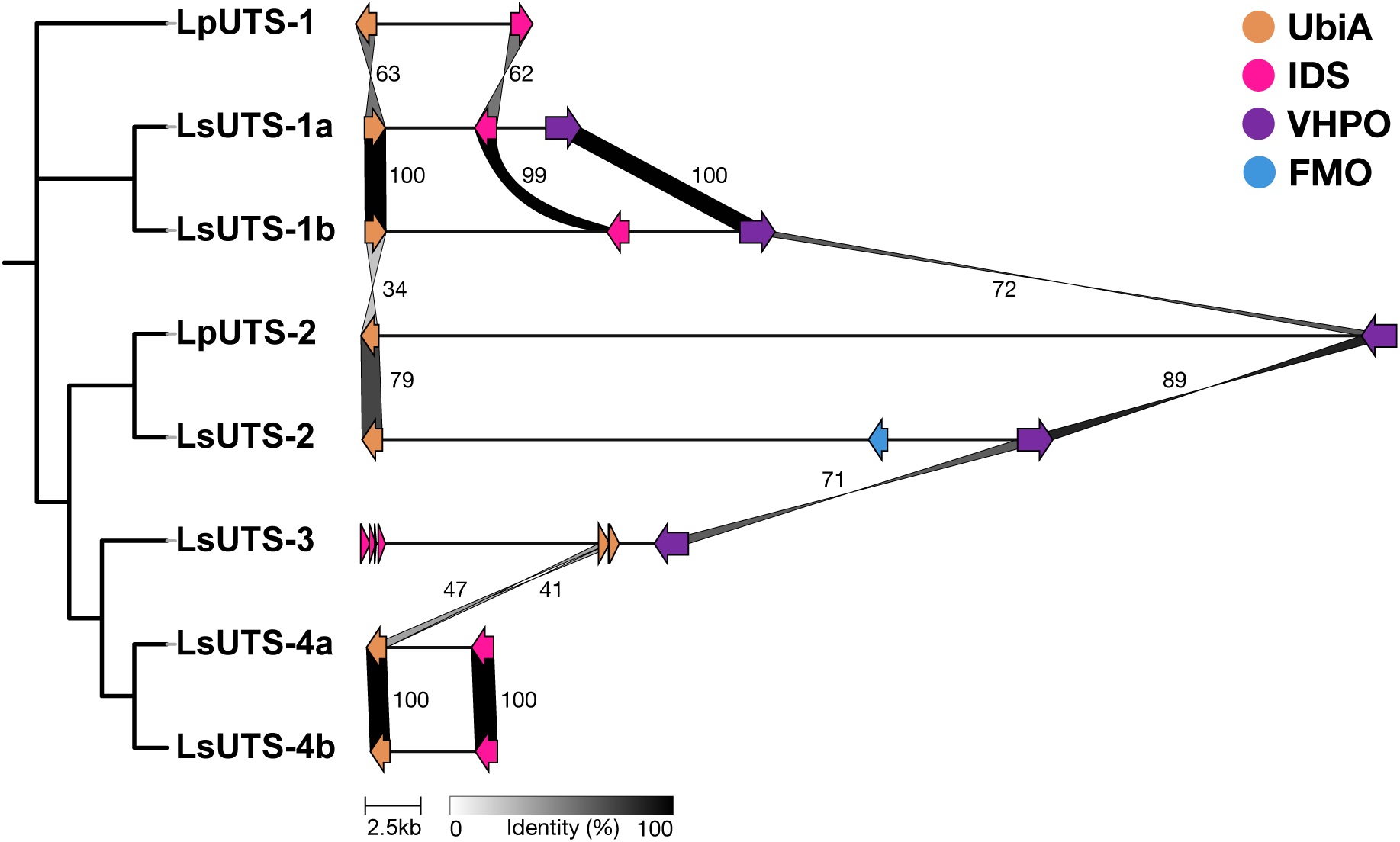
Comparison of eight UbiA-type terpene synthase (UTS) biosynthetic gene clusters identified in *Laurencia* genomes. The phylogenetic tree was generated from UTS sequences aligned using MAFFT, and gene cluster comparisons were visualized using clinker.^16^ Arrows indicate gene positions and orientations and are colored by annotation: UbiA, orange; isoprenyl diphosphate synthase (IDS), pink; vanadium-dependent haloperoxidase (VHPO), purple; and flavin-containing monooxygenase (FMO), blue. Links connect homologous proteins, with shading and numerical labels indicating percentage amino acid identity.

**Figure S7.**
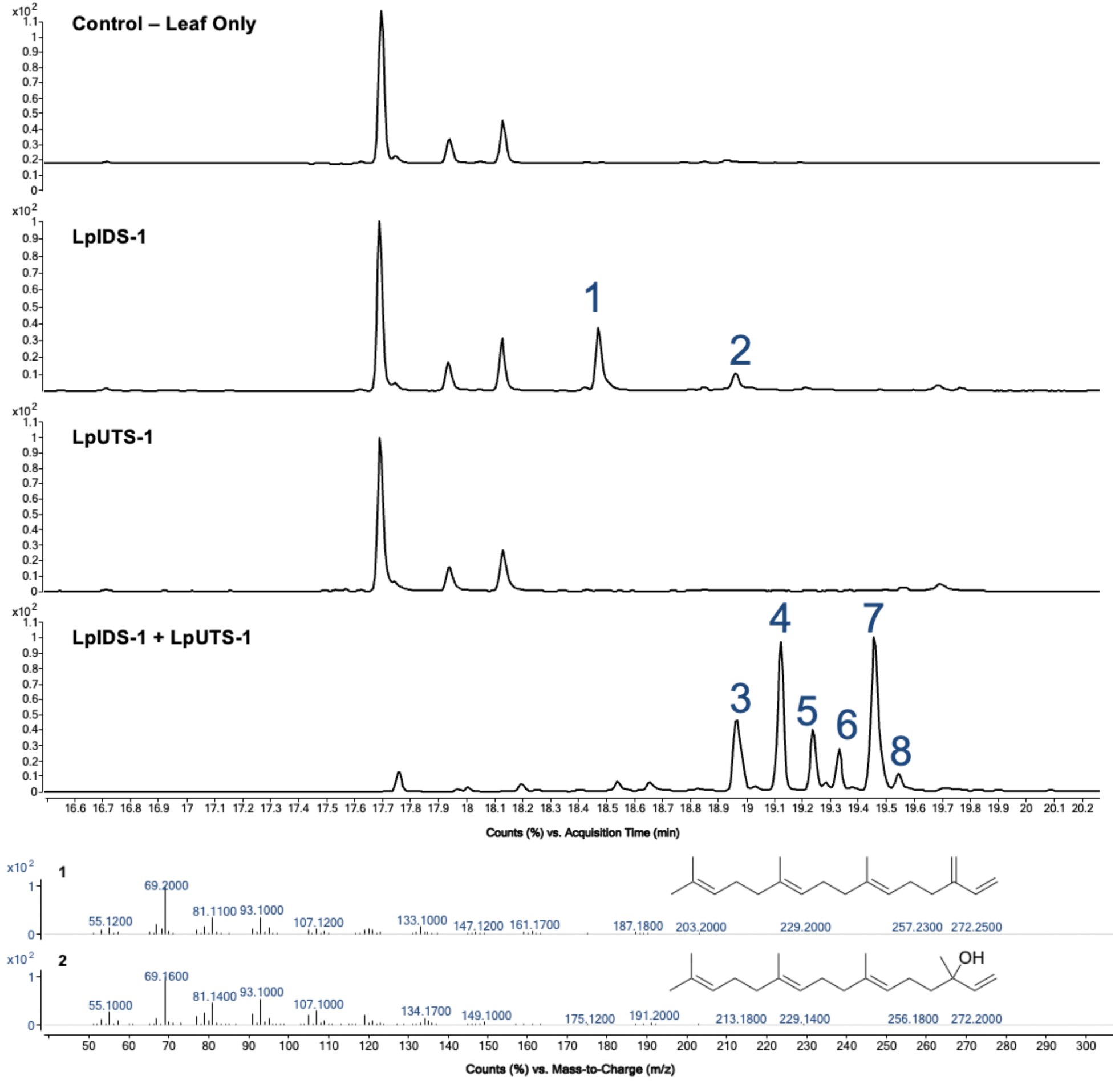
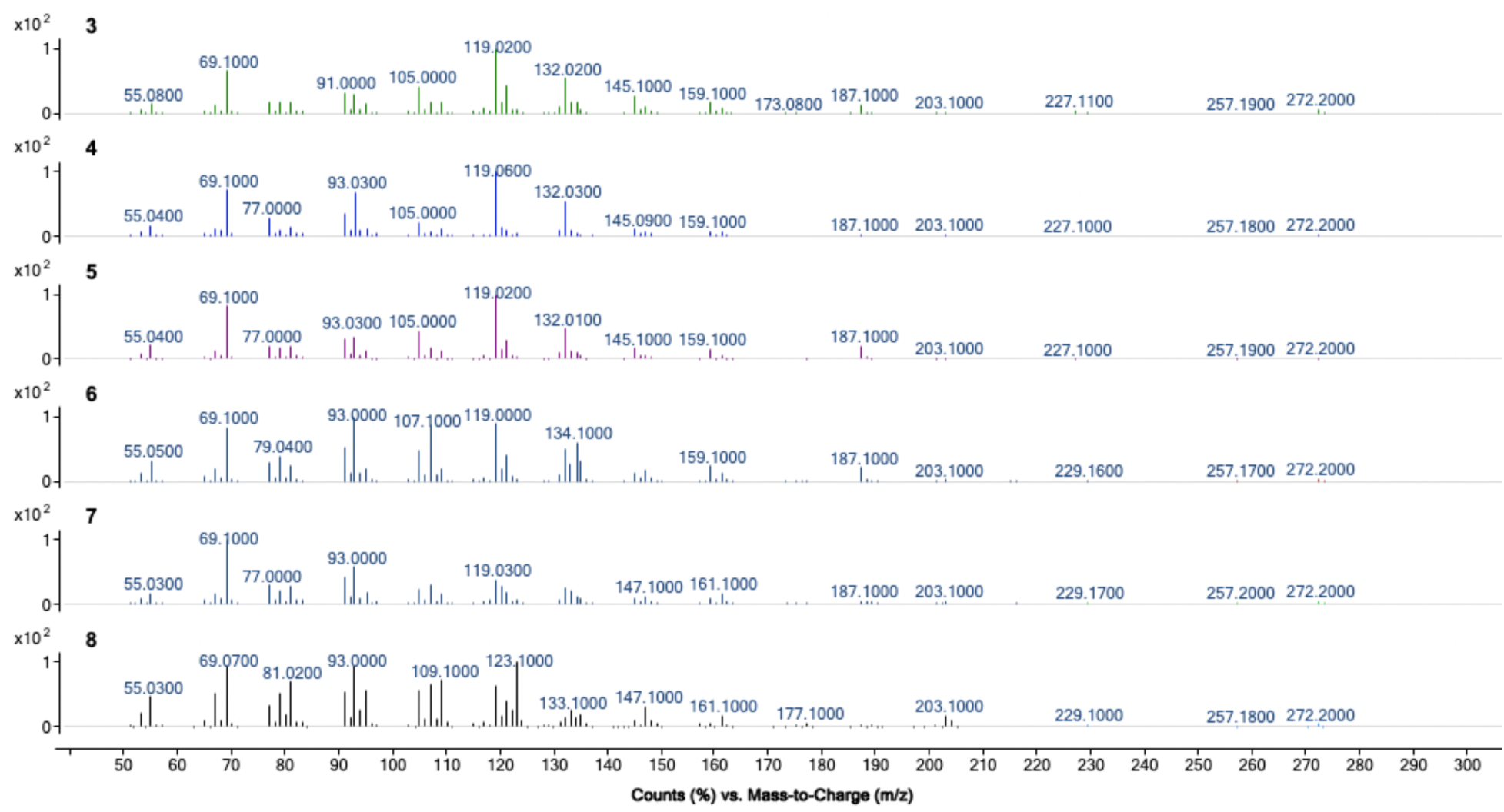
GCMS analysis of LpIDS-1 and LpUTS-1 expressed in *Nicotiana benthamiana*. Total ion chromatograms of extracts from the leaf-only control and leaves expressing LpIDS-1, LpUTS-1, or LpIDS-1 and LpUTS-1 together. Peaks 1–8 indicate metabolites associated with heterologous expression, with the corresponding mass spectra shown below.

**Figure S8.**
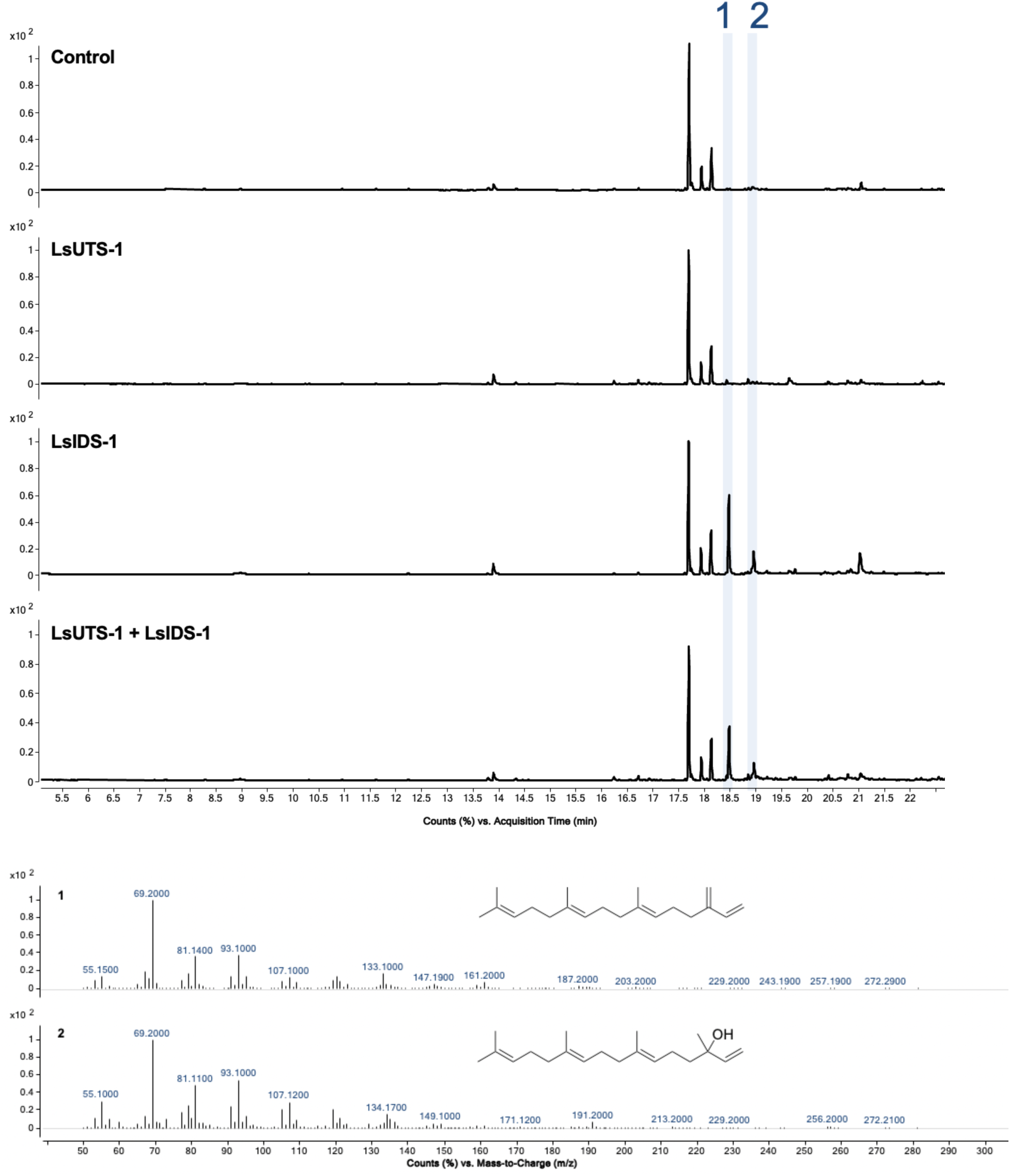
GCMS analysis of LsIDS-1 and LsUTS-1a expressed in *Nicotiana benthamiana*. Total ion chromatograms of extracts from the leaf-only control and leaves expressing LsUTS-1a, LsIDS-1, and LsUTS-1a and LsIDS-1 together. Peaks 1 and 2 indicate metabolites associated with heterologous expression, with the corresponding mass spectra shown below.

**Figure S9.**
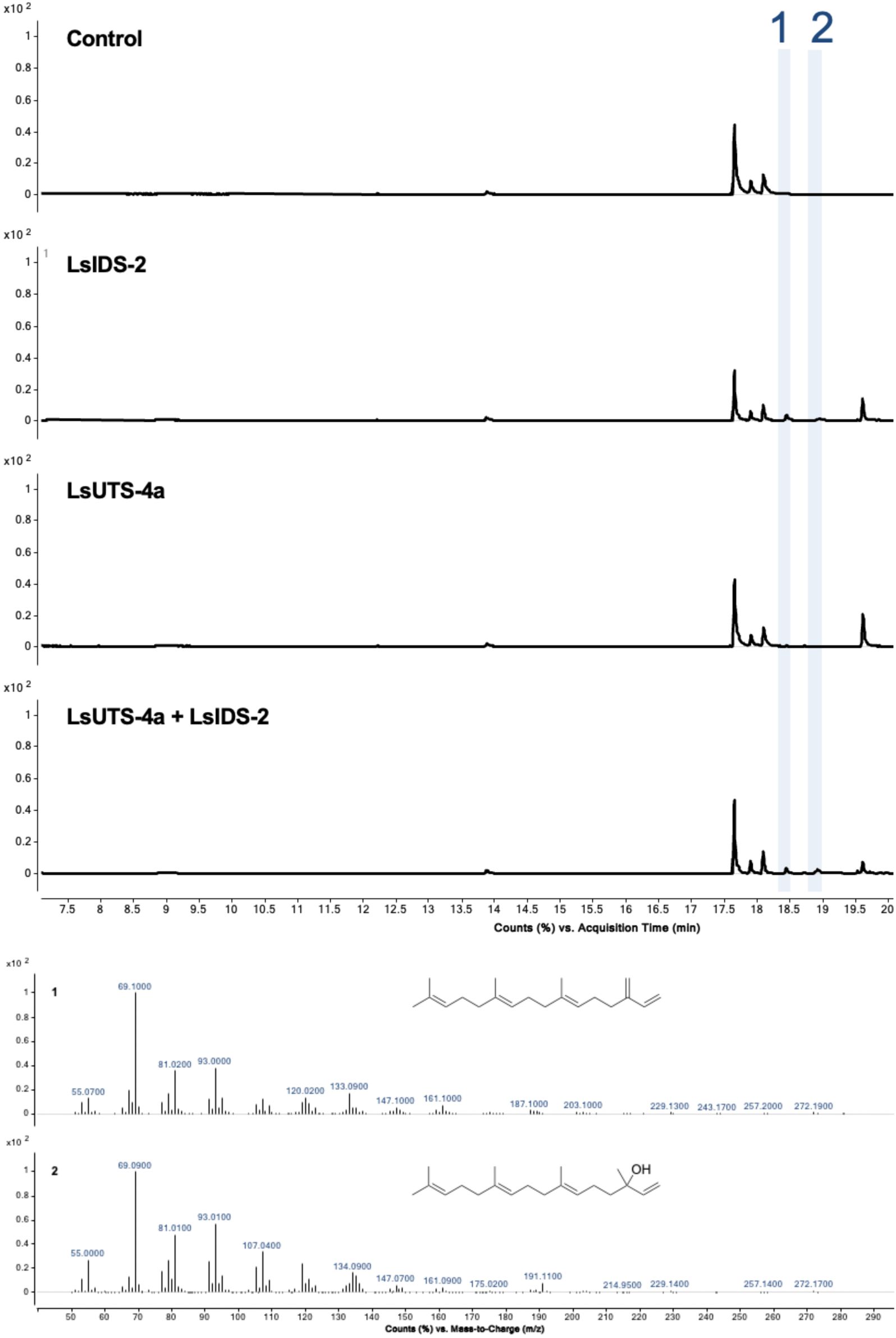
GCMS analysis of LsIDS-2 and LsUTS-4a expressed in *Nicotiana benthamiana*. Total ion chromatograms of extracts from the leaf-only control and leaves expressing LsUTS-4a, LsIDS-2, and LsUTS-4a and LsIDS-2 together. Peaks 1 and 2 indicate metabolites associated with heterologous expression, with the corresponding mass spectra shown below.

**Figure S10.**
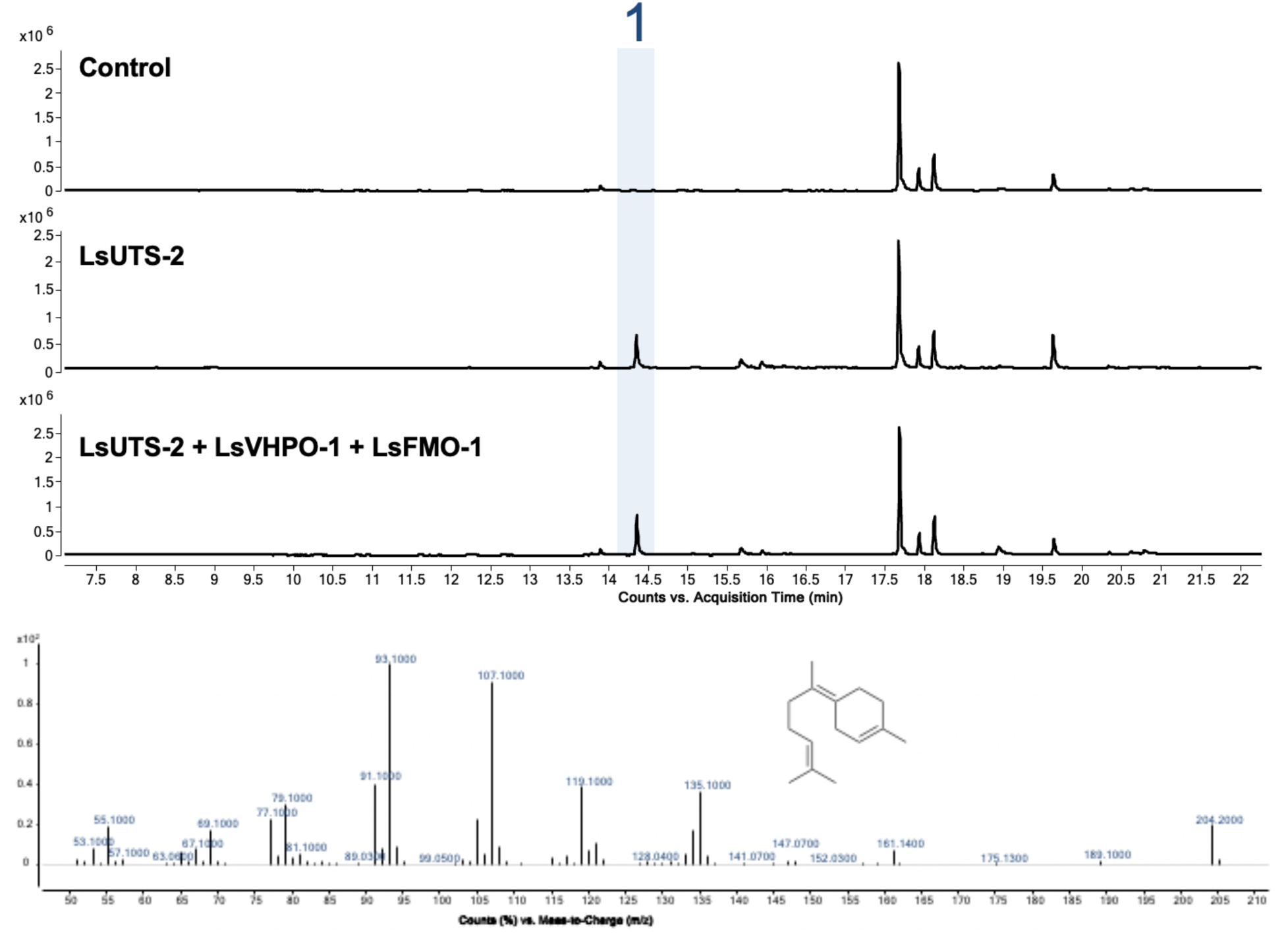
GCMS analysis of LsUTS-2 coexpressed with putative tailoring enzymes. Total ion chromatograms of extracts from the control and samples expressing LsUTS-2 alone or together with LsVHPO-1 and LsFMO-1. Peak 1, detected following expression of LsUTS-2, was identified as (E)-γ-bisabolene based on its mass spectrum, shown below. Coexpression with LsVHPO-1 and LsFMO-1 did not produce additional detectable products under the conditions tested.

**Figure S11.**
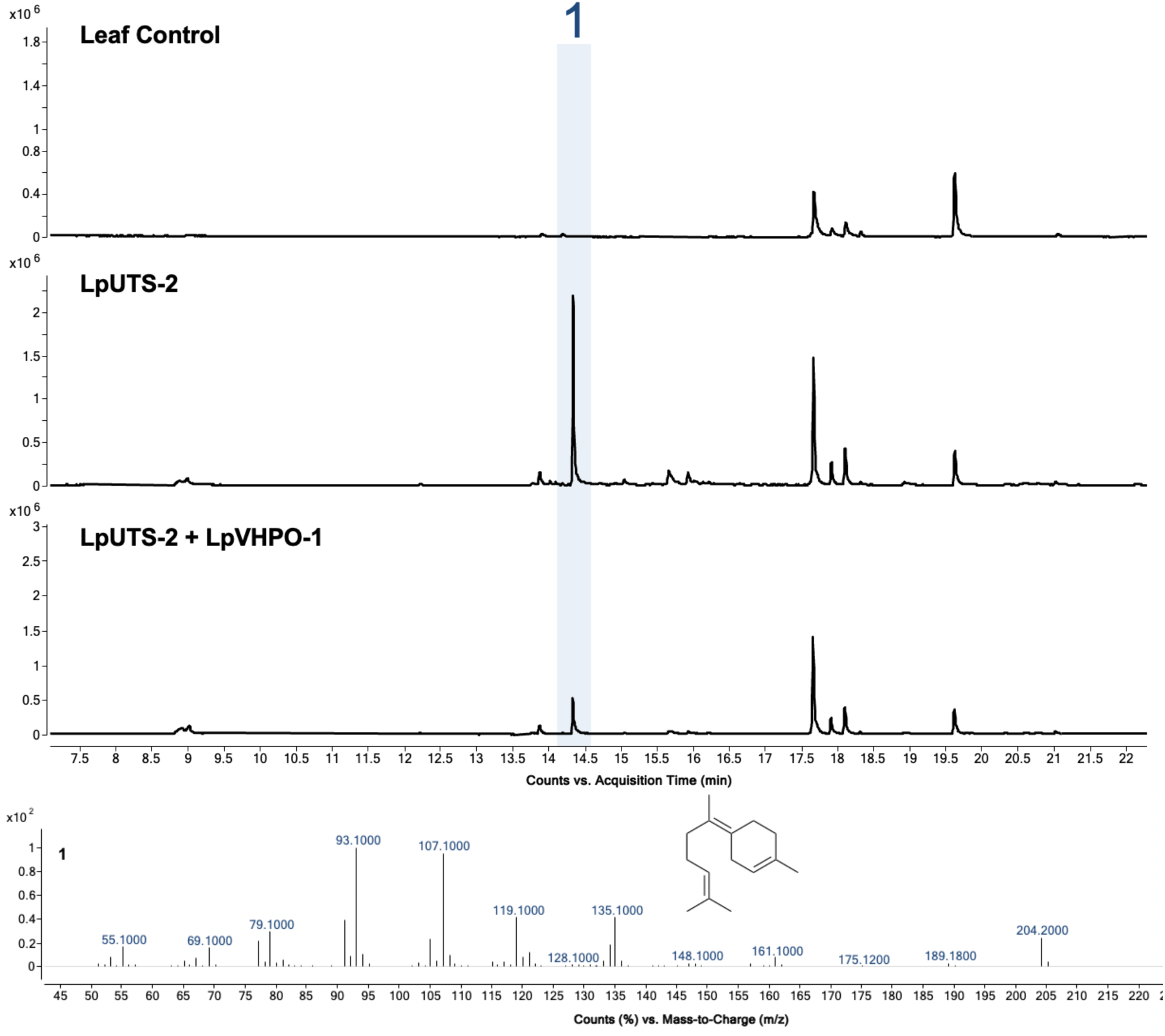
GCMS analysis of LpUTS-2 coexpressed with LpVHPO-1. Total ion chromatograms of extracts from the leaf-only control and *Nicotiana benthamiana* leaves expressing LpUTS-2 alone or together with LpVHPO-1. Peak 1, produced by LpUTS-2, was identified as E-γ-bisabolene based on its electron-ionization mass spectrum, shown below. Coexpression with LpVHPO-1 reduced the abundance of peak 1 but did not yield additional detectable products under the conditions tested.

**Figure S12.**
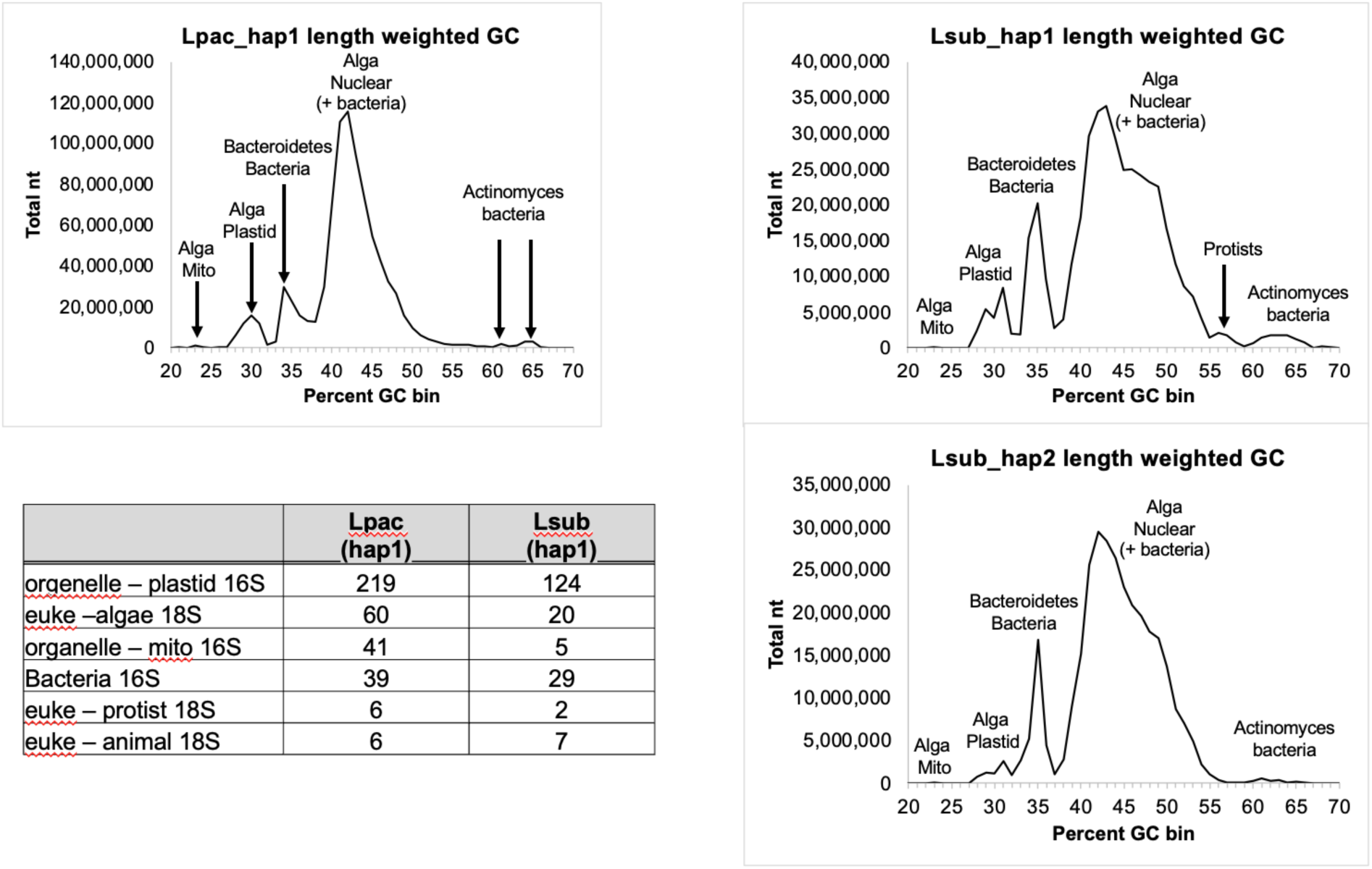
Metagenomic composition of the *Laurencia pacifica* and *Laurencia subopposita* genome assemblies. Length-weighted GC-content distributions are shown for the *L. pacifica* haplotype 1 assembly and the *L. subopposita* haplotype 1 and 2 assemblies. Peaks were assigned to putative algal nuclear, plastid, and mitochondrial sequences or associated bacterial and protist sequences based on their GC content and taxonomic annotation. The table summarizes the numbers of plastid, mitochondrial, bacterial, algal, protist, and animal 16S or 18S rRNA sequences identified in each assembly.

**Figure S13.**
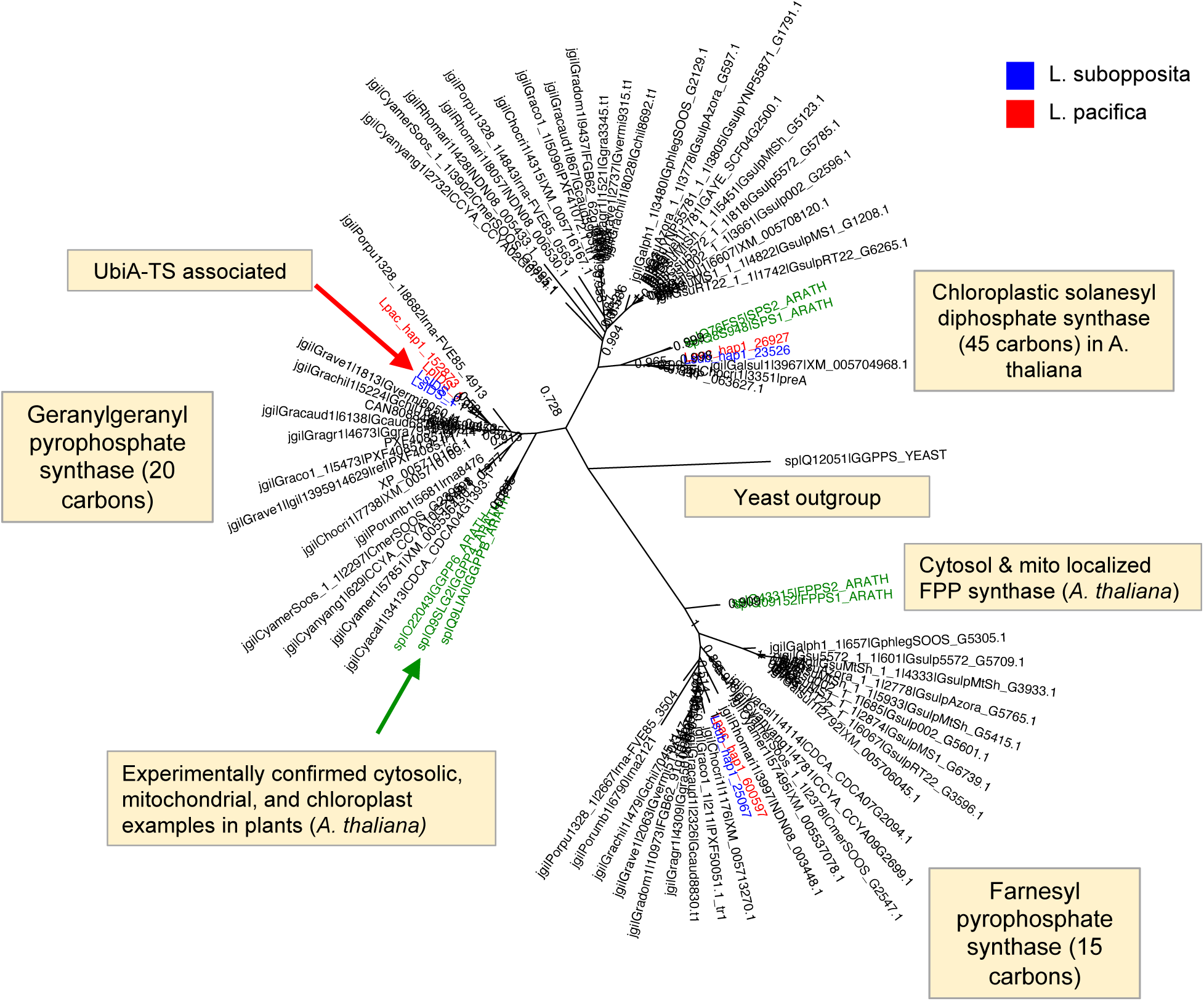
Phylogenetic analysis of *Laurencia* isoprenyl diphosphate synthases (IDSs). Phylogenetic relationships of IDS sequences identified in *L. subopposita* (blue) and *L. pacifica* (red) with characterized and putative IDS homologs. The sequences separate into clades containing geranylgeranyl diphosphate synthases (GGPPSs; C20 products) and farnesyl diphosphate synthases (FPPSs; C15 products). Characterized cytosolic, mitochondrial, and plastid-localized IDSs from *Arabidopsis thaliana* are shown in green as references. The red arrow indicates the *Laurencia* IDS associated with a UbiA-type terpene synthase, and the green arrow indicates experimentally characterized plant IDSs. The yeast GGPPS sequence was used as an outgroup. Selected branch-support values are shown at the nodes.

**Figure S14.**
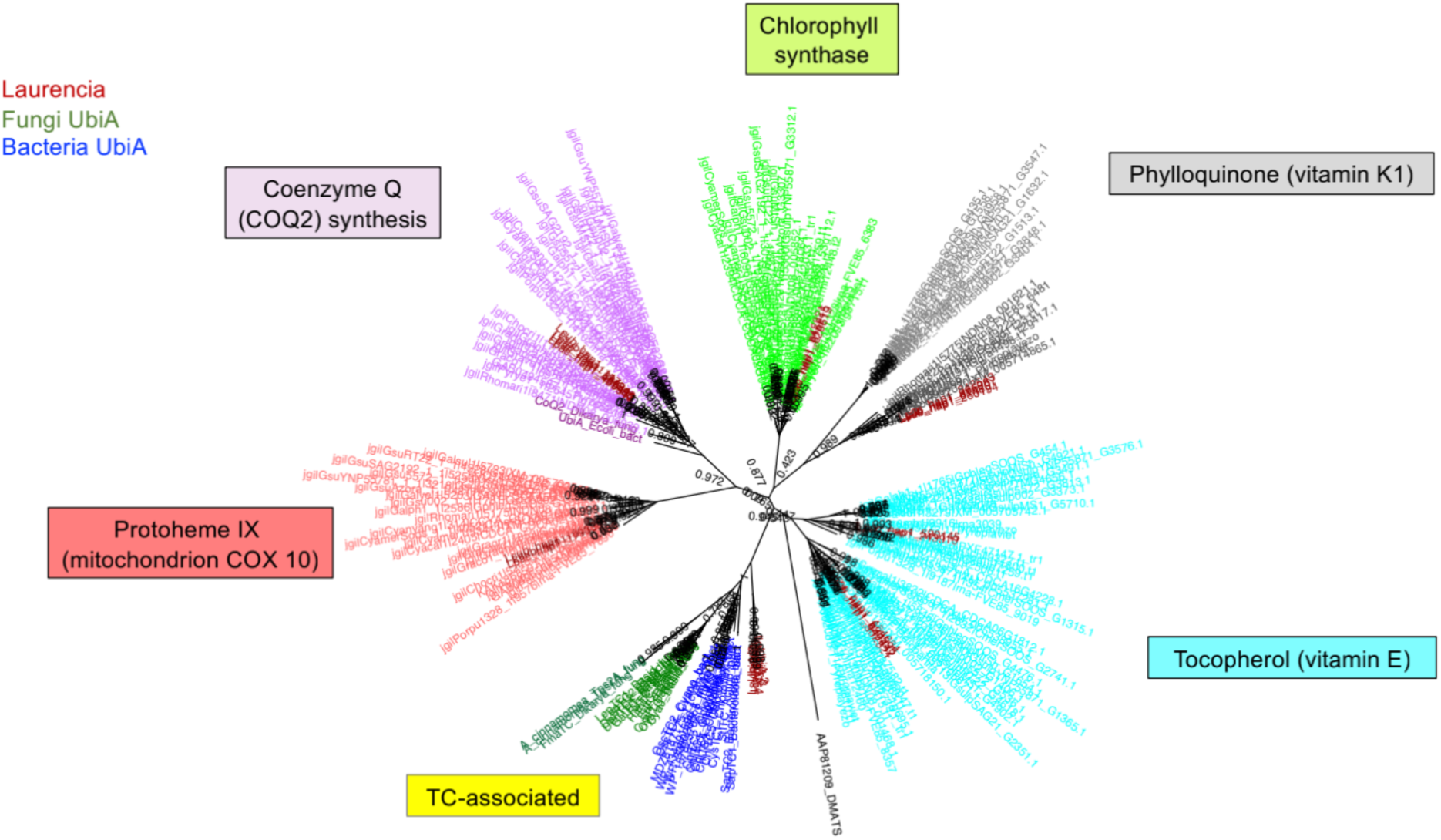
Phylogenetic Analysis of *Laurencia* UbiA prenyltransferases (PTs) and UTSs. Phylogenetic tree showing the relationships between *Laurencia* candidate proteins and reference UbiA-family sequences. Colors distinguish sequence groups, and numbers at internal nodes indicate branch support values. Dimethylallyltryptophan synthase (DMATS) from *Claviceps purpurea* (AAP81209.1) was used as the outgroup.

## Notes

### Competing Interest Statement

The authors have declared no competing interest.

